# Phospho-tuning immunity through Denisovan, modern human and mouse TNFAIP3 gene variants

**DOI:** 10.1101/589507

**Authors:** Nathan W. Zammit, Owen M. Siggs, Paul Gray, Keisuke Horikawa, Stephen R. Daley, David B. Langley, Daniele Cultrone, Elisabeth K. Malle, Stacey N. Walters, Jeanette E. Villanueva, Joanna Warren, Amanda Russell, Mark J. Cowley, Velimir Gayevskiy, Marcel E. Dinger, Claudia Loetsch, Cecile King, Robert Brink, David Zahra, Geeta Chaudhri, Gunasegaran Karupiah, Belinda Whittle, Carla Roots, Edward Bertram, Michiko Yamada, Yogesh Jeelall, Anselm Enders, Benjamin E. Clifton, Peter D. Mabbitt, Colin J. Jackson, Susan R. Watson, Craig N. Jenne, Lewis L. Lanier, Tim Wiltshire, Matthew H. Spitzer, Garry P. Nolan, Frank Schmitz, Alan Aderem, Benjamin T. Porebski, Ashley M. Buckle, Derek W. Abbott, John B. Ziegler, Maria E. Craig, Paul Benitez-Aguirre, Juliana Teo, Melanie Wong, Murray P. Cox, Wilson Phung, Ingrid E. Wertz, Daniel Christ, Christopher C. Goodnow, Shane T. Grey

## Abstract

Resisting or tolerating microbes are alternative strategies to survive infection, but little is known about the evolutionary mechanisms controlling this balance. Here, genomic analyses of anatomically modern humans, extinct Denisovan hominins, and mice revealed a series of missense variants in the immune response inhibitor A20 (encoded by *TNFAIP3*), substituting non-catalytic residues of the ubiquitin protease domain to diminish IκB-dependent phosphorylation and activation of A20. Two A20 variants with partial phosphorylation deficits appeared beneficial: one originating in Denisovans and introgressed in modern humans throughout Oceania, and another in a mouse strain resistant to Coxsackievirus. By contrast, a variant with 95% loss of phosphorylation caused spontaneous inflammatory disease in humans and mice. Analysis of the partial phosphorylation variant in mice revealed diminished tolerance of bacterial lipopolysaccharide or to poxvirus inoculation as trade-offs for enhanced immunity.

**One Sentence Summary:** Modern and ancient variants reveal a genetically tunable element for balancing immunity and microbial tolerance.

## Main Text

Microbial resistance involves innate and adaptive immune responses that prevent, diminish, or clear infection, often causing collateral damage to host tissues and increased energy demands. Studies in plants^1^, and more recently in animals^2,3^, have shown that in some circumstances it can be more efficient for a host to tolerate microbes rather than resist them. Microbial tolerance involves homeostatic mechanisms to raise thresholds for initiating immune responses, to physically separate microbes from host immune receptors, and to repair damage caused directly by microbes or by collateral inflammation^3-6^. The genetic means by which microbial resistance and tolerance are balanced remains incompletely understood. Population genetic modelling predicts that resistance traits favor host polymorphism and microbial evasion, whereas microbial tolerance traits tend towards fixation in hosts and microbial mutualism^7^. High mortality in indigenous human populations of Oceania and the Americas exposed to smallpox demonstrate how a tolerated pathogen in its adapted host can cause devastating disease when introduced into non-adapted populations^8,9^. An example in animals can be seen for European rabbits exposed to myxoma poxvirus originating from South American rabbits^8^. These cases illustrate the importance of fine-tuning microbial immunity and tolerance during co-evolution of host and microbe, although little is known about the molecular pathways which underpin it.

A binary perspective of the importance of balancing microbial immunity and tolerance comes from Mendelian gene variants in mice and humans that completely inactivate the function of one or both alleles of genes such as *CTLA4, IL10*, *FOXP3*, or *TNFAIP3*^10-12^. These cause severe pediatric autoimmune or inflammatory disease, particularly at mucosal barriers where large microbial populations are normally tolerated, such as the microbial burden that drives inflammatory pathology in mouse *Tnfaip3* deficiency^6^. Genome-wide and candidate association studies in humans implicate variants at or near the *TNFAIP3* locus with susceptibility to autoimmune disease^13^. In contrast to these disease-associated traits, few examples of beneficial genetic adjustments that decrease microbial tolerance in favor for heightened immunity exist.

A20, encoded by the *TNFAIP3* gene, promotes microbial tolerance as a negative regulator of NF-κB signaling: an evolutionarily ancient and central pathway for activating innate and adaptive immune responses^13^. A20 has multiple domains with inhibitory activity against NF-κB, primarily preventing activation of the central IκB kinase (IKK) by upstream proteins RIPK1, TRAF6 and NEMO. The A20 ovarian tumor (OTU) domain is a deubiquitin (DUB) protease that cleaves activating K63-linked ubiquitin chains from RIPKI, TRAF6 or NEMO^14-16^. The A20 zinc finger 7 domain (ZnF7) binds linear polyubiquitin to suppress IKK activation, whereas ZnF4 promotes ligation of K48-linked ubiquitin chains to RIPK1, triggering RIPK1 proteolysis^14,17^. A20 feedback inhibition is induced at two levels: NF-κB proteins directly induce *TNFAIP3* mRNA, and the inhibitory activities of A20 protein are enhanced by IκKβ-induced serine phosphorylation near the ZnF domains, notably S381^18,19^.

The role of the A20 OTU domain nevertheless remains enigmatic. The ZnF domains alone are sufficient for NF-κB inhibitory function in cell-based studies^20,21^, and mice homozygous for *TNFAIP3* missense variants creating a catalytically inactive OTU domain have little^15,18^ or no evidence^20^ of excessive NF-κB signaling. Here we demonstrate that anatomically modern human, archaic Denisovan, and mouse missense variants in the OTU domain modulate A20 phosphorylation by IKK, serving as a genetically tunable element with profound effects on the balance between microbial tolerance and immunity, and evidence of introgression to high frequencies during human history.

## Modern human A20 OTU variants acquired from Denisovans

Three convergent sequencing studies led us to a unique functional class of A20 missense variants affecting the OTU domain, separate from the ubiquitin protease catalytic site (Fig. 1A, B; Extended Data Fig. 1). The most subtle allele, comprising T108A (rs376205580 g.138196008A>G) and I207L (rs141807543 g.138196957A>C) missense variants in *cis*, was identified by whole genome sequencing in four of 85 families in Sydney. The majority of individuals in our cohort carrying the T108A;I207L allele were healthy family members of Māori or Pacific Islander ancestry. These variants were rare in a public variant collection (gnomAD r2.0.2) but most frequent among individuals with an unassigned ancestry (Extended Data Fig. 2A).

**Fig. 1.**
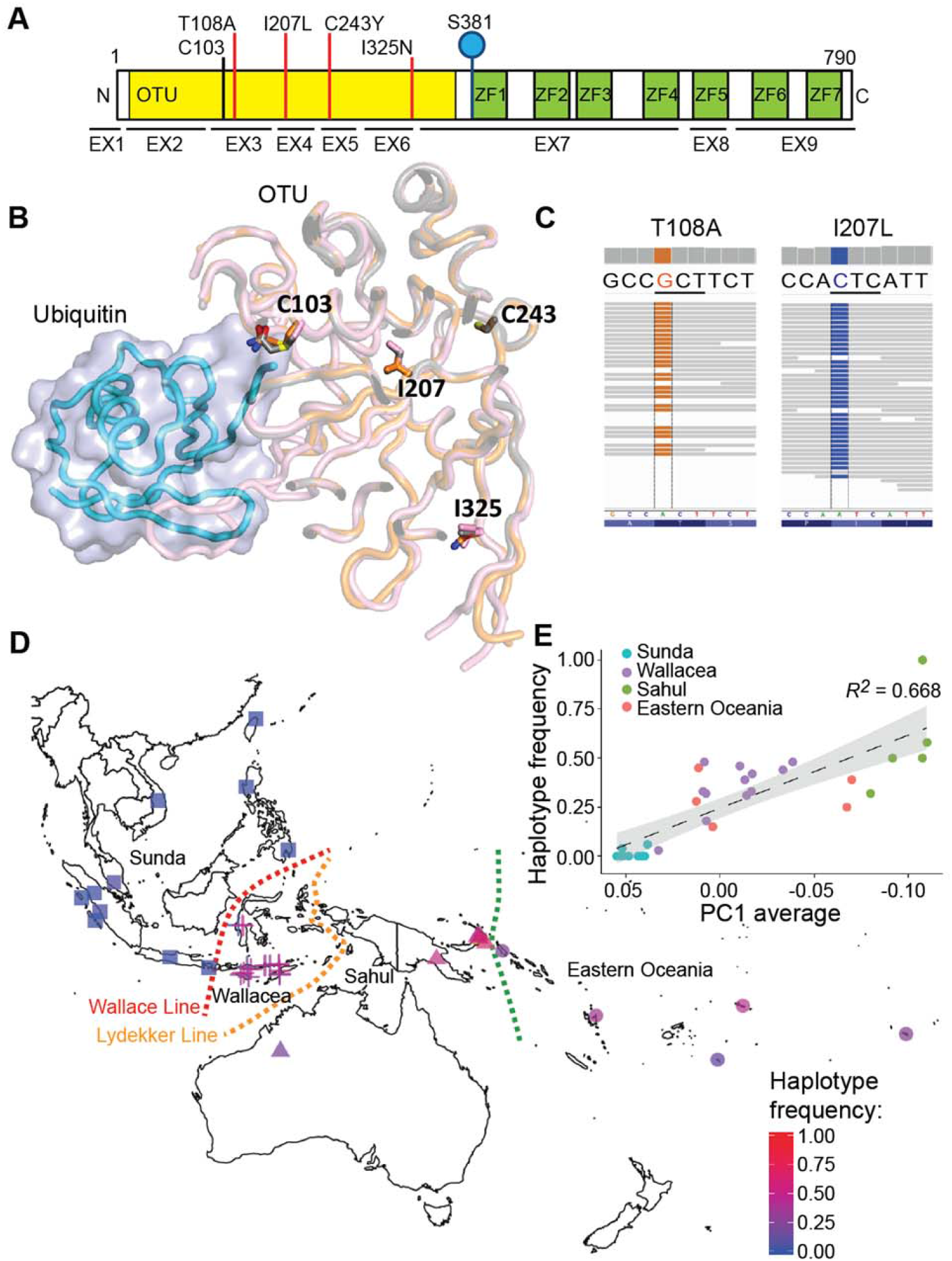
Denisovan, modern human, and mouse variants substituting noncatalytic residues of the A20 OTU domain. **(A)** Schematic of A20 protein encoded by the *TNFAIP3* gene (CCDS5187), showing exons (EX1-9), ovarian tumor domain (OTU), zinc fingers (ZF1- 7), OTU catalytic residue (C103), S381 phosphorylation site, and missense variants studied here (red lines). **(B)** Location of variant residues in sliced structure 5LRX^42^ of A20 OTU domain (pink) complexed with ubiquitin (blue). Structures for WT and I325N OTU domains are superposed in grey and orange, respectively. **(C)** Read data of a high-coverage Denisovan genome^27^ across *TNFAIP3* codons 108 and 207. **(D)** Imputed frequency of the Denisovan *TNFAIP3* haplotype in genotype array data from 514 individuals from indigenous populations across Island Southeast Asia and Oceania^30^, and directly observed in exome data from 72 Martu indigenous Australians^32^. Different shape symbols group populations from Sunda, Wallacea, Sahul and Eastern Oceania. **(E)** Imputed Denisovan *TNFAIP3* haplotype frequency from populations described in (D) (filled circles) versus a surrogate estimate of Papuan ancestry (principal component 1, Extended Data Fig. 3). Gray shading indicates 95% confidence interval of linear regression line.

We traced the global distribution of the T108A;I207L allele in 279 individuals from the Simons Genome Diversity Project dataset^22^, revealing high frequencies ranging from 25-75% among people of Island Southeast Asia and Oceania, but an absence of the allele elsewhere in the world (Extended Data Fig. 2B, Extended Data Table 1).

Unlike Africans or Eurasians, people in Island Southeast Asia and Oceania acquired up to 5% of their genome from Denisovans: archaic hominins that interbred with modern humans ∼50,000 years ago migrating through Asia to settle the continent of Sahul (now Papua New Guinea and Australia)^23-26^. Analysis of the high-coverage genome of a Denisovan finger phalanx from a cave in the Altai Mountains of Siberia^27^ revealed homozygous T108A;I207L variants (Fig. 1C). Both variants were absent from the genome of a Neanderthal who had inhabited the same cave^28^ (Extended Data Table 2), suggesting that T108A and I207L arose after the divergence of Denisovan and Neanderthal lineages 170,000-700,000 years ago^27^.

Multiple Denisovan-derived genomic regions, including one encompassing *TNFAIP3*, bear strong signatures of introgression in Papuans^25,26,29^. By imputing haplotypes (Extended Data Table 2) using genotype array data from 514 individuals from indigenous populations across Island Southeast Asia and Oceania^30^, we found evidence that the Denisovan *TNFAIP3* haplotype has reached high frequencies in populations east of the Wallace Line, a 50 million year-old faunal boundary separating organisms of Asiatic and Australian origin via deep water channels between the two continental shelves (Fig. 1D, E and Extended Data Fig. 3)^31^. Frequencies ranged from 0% in almost all populations west of the Wallace Line, but reached 100% in 8 Baining individuals from eastern Papua New Guinea. We observed the same haplotype in 31/144 (22%) of exome-sequenced alleles in Martu Indigenous Australians from the Pilbara region of Western Australia^32^, implying the haplotype enrichment occurred before the isolation of Indigenous Australian and Papuan populations. The high frequency in Polynesia indicates the Denisovan *TNFAIP3* haplotype was retained after admixture with Austronesian farming populations expanding from mainland Asia starting ∼4,000 years ago^33^ in Eastern Indonesia and ∼3,000 years ago in the Southwest Pacific^34^.

Aside from the two missense variants in the introgressed Denisovan *TNFAIP3* haplotype, many other non-coding variants could conceivably modulate transcription. Deep sequencing of cDNA nevertheless revealed equal amounts of *TNFAIP3* mRNA from Denisovan and modern human alleles in heterozygous leukocytes, with or without TNFα stimulation (Extended Data Fig. 4). Of the two coding variants, T108A was not predicted computationally to alter A20 function (Phred-scaled CADD score of 0.002) and occurs in other vertebrate species (Extended Data Fig. 1). By contrast, I207 is invariant across most jawed vertebrates, and I207L was predicted to be the most deleterious variant across the Denisovan *TNFAIP3* haplotype (Phred-scaled CADD score: 23.2; Extended Data Fig. 3E).

## A mouse A20 OTU variant confers heightened resistance to Coxsackievirus

The results above identify an ancient substitution in the A20 OTU domain as apparently beneficial in human history. Further evidence that A20 OTU domain variants could be beneficial came from another OTU substitution, I325N, identified in a genome-wide mouse mutagenesis screen segregating with increased frequencies of circulating CD44^hi^ activated/memory T cells and regulatory T cells (Fig. 2A) in otherwise healthy adult mice. Detailed analysis revealed the I325N mutation also increased numbers of B cells in the spleen and peritoneal cavity, and diminished IκBα within B cells, CD8 T cells, NK cells and dendritic cells (Fig. 2B and Extended Data Fig. 5). Macrophages immortalized from bone marrow of I325N mutant mice produced more inflammatory cytokines in response to lipopolysaccharide (LPS) than controls from wild-type mice (Extended Data Fig. 6). In thymocytes, I325N increased TNFα-induced NF-κB signaling in ways consistent with diminished A20-mediated inhibition (Extended Data Fig. 7)^18^.

**Fig. 2.**
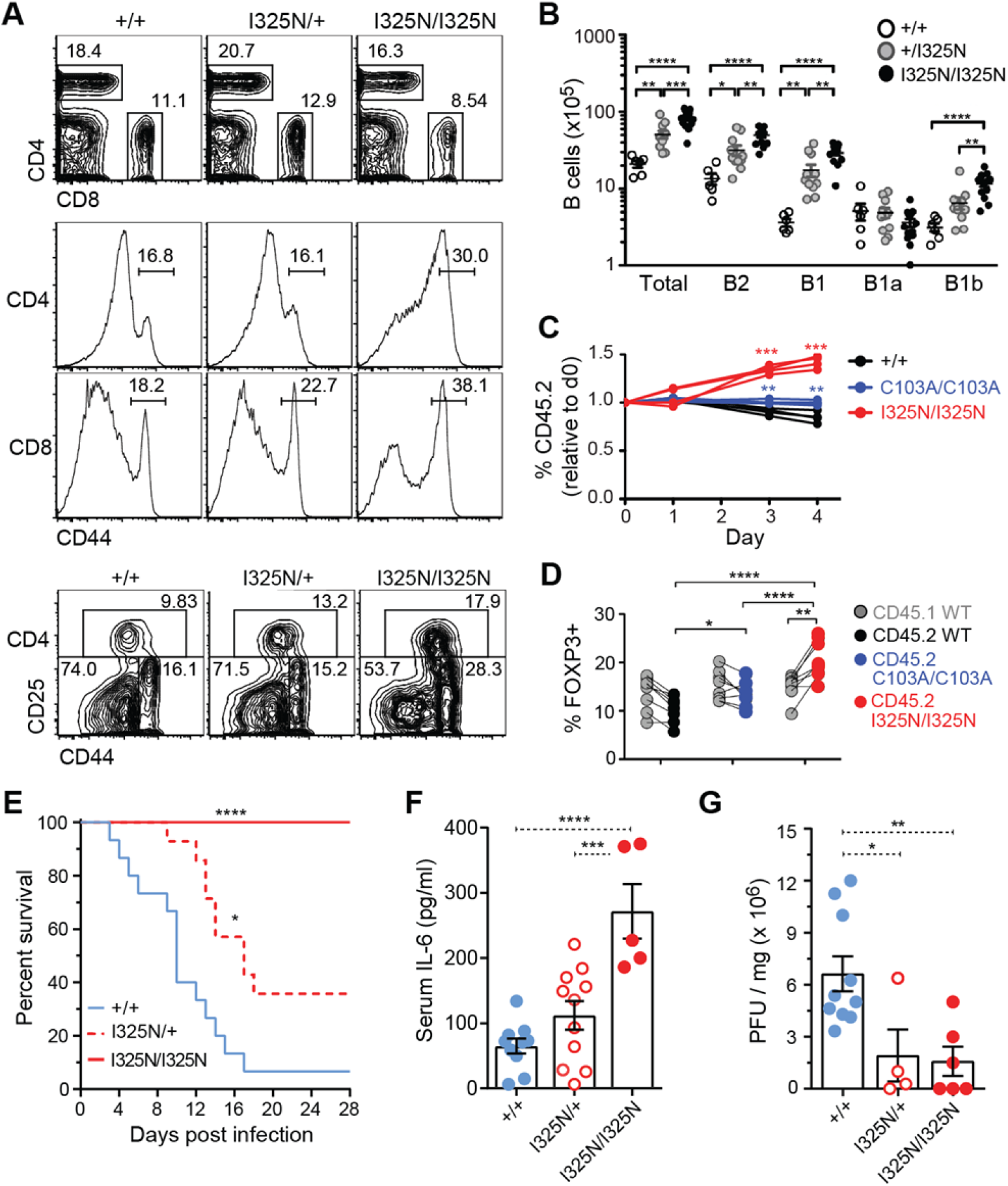
I325N OTU variant confers heightened immunity in mice. **(A)** Representative flow cytometric analysis of splenocytes, showing percentage of CD4^+^ or CD8^+^ T cells, and percentage CD44^hi^ activated/memory cells within these subsets or percentage CD25^+^ CD44^int^ regulatory T- cells or CD25^-^ CD44^high^ effector/memory T cells among CD4^+^ cells. **(B)** Numbers of indicated B cells in peritoneum of individual *Tnfaip3* mice (circles). **(C, D)** Wild-type CD45.1^+^ mice transplanted with equal mixtures of congenic bone marrow from *Tnfaip3*^+/+^ CD45.1^+^ donors and CD45.2^+^ donors of *Tnfaip3*^I325N/I325N^, *Tnfaip3*^C103A/C103A^ or *Tnfaip3*^+/+^ genotypes. (C) Splenocytes from chimeras cultured with 0.1 µg/ml LPS for 0-4 days, and the frequency of CD45.2^+^ cells of the indicated *Tnfaip3* genotypes was measured among B cells from individual chimeras (lines), expressed relative to their frequency on day 0. (D) Pairwise comparison of frequency of FOXP3^+^ cells amongst CD45.2^+^ and CD45.1^+^ CD4^+^ splenocytes from the same chimera. **(E-G)** Female *Tnfaip3*^+/+^ (*n*=15), *Tnfaip*3^I325N/+^ (*n*=14) or *Tnfaip*^3I325N/I325N^ (*n*=10) C57BL/6 mice were injected with Coxsackie B4 virus strain E2 on day 0. (E) Kaplan-Meier survival data (Log-rank test). (F) Plasma IL-6 and (G) virus titers (plaque forming units, PFU) per mg of pancreas on day 3. Error bars represent SEM and one-way ANOVA used for significance analysis unless otherwise stated, \**P* < 0.05; \*\**P* < 0.01; \*\*\**P* < 0.001; \*\*\*\**P* < 0.0001.

In wild-type mice transplanted with mixtures of mutant and wild-type A20 bone marrow, the I325N mutation acted cell autonomously to increase B cell activation and proliferation by LPS or antigen receptors, and to increase TCR and CD28-dependent formation of FOXP3^+^ CD4^+^ regulatory T cells and their Helios^+^ FOXP3^-^ precursors within the thymus (Fig. 2C, D, Extended Data Fig. 8). Surprisingly, I325N had a greater effect than the C103A OTU domain mutation analyzed in parallel bone marrow transplant recipients, despite C103A completely inactivating the polyubiquitin protease activity of A20^15,20^, indicating I325N must diminish additional inhibitory mechanisms.

Consistent with heightened cellular markers of immunity, I325N mutant mice had greater resistance to Coxsackievirus B4 strain E2, a virus isolated from a human neonate with a disseminated fatal infection causing extensive pancreatic necrosis^35,36^. A virus dose that was lethal for 90% of wild-type C57BL/6 mice was not lethal for *Tnfaip3*^I325N/I325N^ littermates, and caused less mortality in *Tnfaip3*^I325N/+^ mice (Fig. 2E; Extended Data Fig. 9A). Mutant mice had less infectious virus and viral RNA in the pancreas, less mRNA encoding immune response cytokines IL-1β and IFNβ, less pancreatic necrosis, higher serum IL-6, and preserved body-weight and euglycemia (Fig. 2F, G; Extended Data Fig. 9B-G).

The homogeneous genetic background of I325N mutant mice allowed testing if heightened immunity imposed a subclinical cost or altered the insulin anabolic axis^37^*. Tnfaip3*^I325N/+^ mice were healthy, of normal weight, and fertile, producing homozygous mutant offspring at the expected ratio. Homozygotes also appeared healthy, although their body weight was 5-20% less than heterozygous or wild-type littermates (Fig. 3A), with histological analysis revealing low-grade inflammation of the pancreatic islets, colon, kidney, and liver (Fig. 3B-C, Extended Data Fig. 10). Pancreatic insulitis in I325N homozygotes was associated with a 50% reduction in beta cell mass (Fig. 3D), although random blood glucose levels and glucose tolerance tests were normal (Extended Data Fig. 11A-E). Isolated *Tnfaip3*^I325N/I325N^ islets exhibited normal basal insulin output but reduced insulin secretion when stimulated *in vitro* (Extended Data Fig. 11F). Islet transplant and culture experiments showed the A20 mutation acted within islet cells, exaggerating canonical and non-canonical NF-κB signaling, lowering insulin secretion, and increasing inflammatory cytokine production (Fig. 3E-H and Extended Data Fig. 11G-I & 12)^38^.

**Fig. 3.**
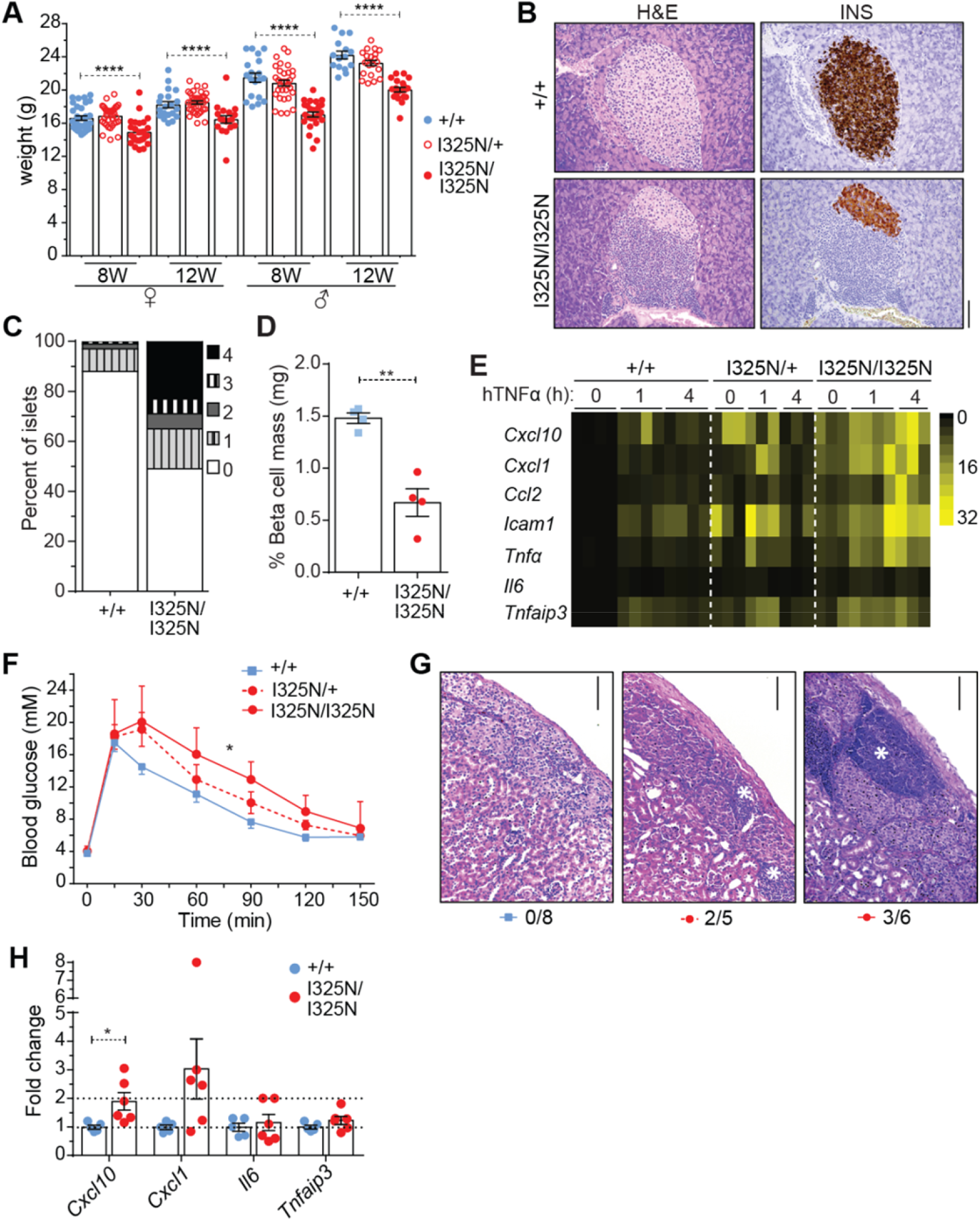
Subclinical inflammatory and metabolic consequences of the I325N variant. **(A)** Body weights of *Tnfaip3*^+/+^, *Tnfaip3*^I325N/+^ *Tnfaip3*^I325N/I325N^ mice of the indicated age and sex. Statistical analysis by unpaired one-way ANOVA: \*\*\*\**P* < 0.0001. **(B)** Hematoxylin and eosin (H&E) and insulin (INS) stained pancreas sections (scale = 100 µm) with **(C)** cumulative insulitis scores (4 represents >75% islet mononuclear cell infiltration, and 0 an absence of infiltrating cells) and **(D)** calculated beta cell mass for *Tnfaip3*^+/+^ and *Tnfaip3*^I325N/I325N^ mice. **(E)** Expression of TNFα-induced genes in islets from *Tnfaip3*^+/+^, *Tnfaip3*^I325N/+^ *Tnfaip3*^I325N/I325N^ mice. Data represents 3 independent islet preparations with 3- 4 biological replicates. **(F)** Glucose tolerance test in diabetic wild-type mice transplanted with *Tnfaip3*^+/+^, *Tnfaip3*^I325N/+^ or *Tnfaip3*^I325N/I325N^ islets, N>8 mice per group. **(G)** H&E sections of islet grafts at post-operative day 30 (scale = 50 µm) with the fraction of grafts exhibiting immune infiltrate shown below. **(H)** Islet grafts isolated on post-operative day 10 were analysed for indicated mRNAs by RT-qPCR. *P* values represent Student’s *t*-test unless otherwise stated, \**P* < 0.05; \*\**P* < 0.01; \*\*\*\**P* < 0.0001.

## An allelic series of OTU domain variants with graded reductions of A20 phosphorylation

The apparent beneficial effects of the T108A;I207L and I325N OTU domain substitutions above contrasted with a third OTU domain substitution, C243Y, found as a family-specific variant causing dominant Mendelian inflammation resembling Behçet’s disease, with childhood-onset oral and genital ulceration and skin inflammation^39^. The biochemical basis for the clinically penetrant effects of C243Y was obscure, because other similarly affected cases of haploinsufficiency of A20 (HA20) result from nonsense or frameshift variants that truncate or eliminate A20 protein^12^.

Despite different clinical consequences, the three OTU domain variants formed a graded biochemical allelic series when full-length A20 and IKKβ were co-expressed in mouse insulinoma cells and S381-phosphorylated A20 was compared with total A20 protein by Western blotting. None of the variants affected A20 protein accumulation but each affected IKKβ-mediated S381 phosphorylation (Fig. 4A-C; Extended Data Fig. 13A). The pathogenic C243Y variant caused the most severe loss of function, diminishing IKKβ-mediated S381 phosphorylation to 5% of wild-type, whereas the Denisovan T108A;I207L variant diminished phosphorylation the least (80% of wild-type) and the non-pathogenic I325N variant had an intermediate effect (50% of wild-type). By contrast, the C103A catalytic site substitution did not cause a significant decrease in A20 phosphorylation. Loss of S381 phosphorylation was also observed by mass spectrometry of A20 purified from unstimulated human cells transfected with A20^I325N^ compared to A20^WT^, whereas A20^C103A^ exhibited normal phosphorylation (Extended Data Fig. 13B).

**Fig. 4.**
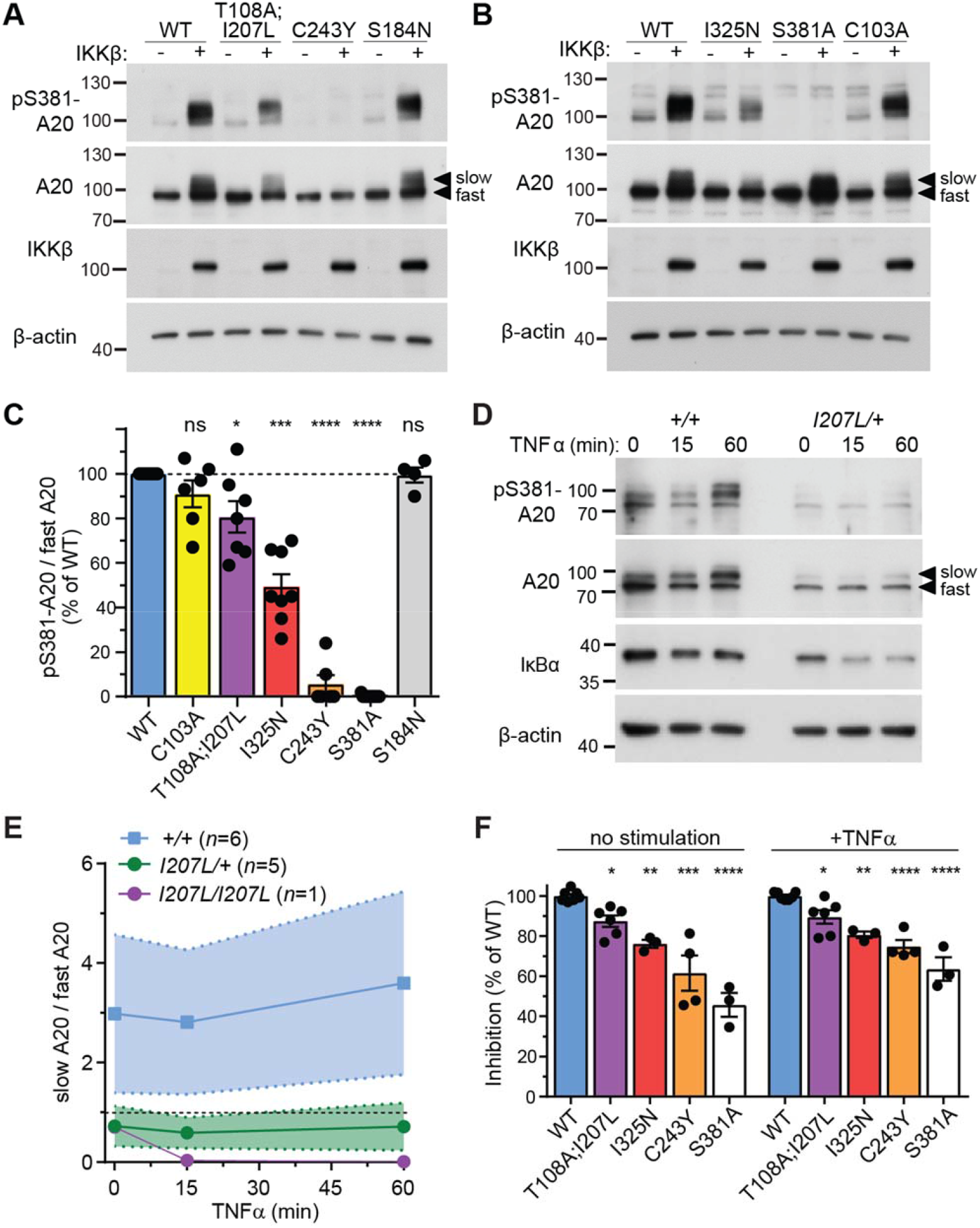
OTU domain variants decrease A20 phosphorylation and NF-κB inhibition. **(A-C)** Representative western blot of lysates from βTC3 mouse insulinoma cells transfected with vectors encoding wild-type (WT) A20 or the indicated A20 mutants, with or without co-transfected IKKβ, and probed with antibodies to phospho-Ser381 A20, total A20, IKKβ and β-actin. Molecular weight markers (kD) and slow-or fast-migrating A20 species are indicated. **(C)** Densitometric analysis from multiple independent experiments (dots) showing the ratio of pSer381-A20 to total fast migrating A20 in IKKβ-cotransfected cells, expressed relative to the ratio in cells with WT A20. **(D)** Representative western blot, probed with antibodies to the indicated proteins, of lysates from peripheral blood mononuclear cells from healthy donors with wild-type A20 or heterozygous for the Denisovan T108A;I207L haplotype, stimulated with TNFα for the indicated times. **(E)** Densitometric ratio of slow-to fast-migrating A20 species, showing mean and standard deviation from six normal donors, five T108A;I207L heterozygotes, and one T108A;I207L homozygote (Extended Data Fig. 14). **(F)** Inhibition of an NF-κB luciferase reporter in βTC3 cells by co-transfection of WT A20 or the indicated A20 mutants, expressed relative to wild-type A20 (Extended Data Fig. 13). Statistical analysis in C and F by one-way ANOVA: ns, *P* > 0.05; \**P* < 0.05; \*\**P* < 0.01; \*\*\**P* < 0.001; \*\*\*\**P* < 0.0001.

S381-phosphorylated A20 predominantly migrated more slowly in SDS-PAGE than unphosphorylated A20, either when tested by co-transfection with IKKβ (Fig. 4A, B) or in TNFα-stimulated peripheral blood leukocytes with endogenous A20 and IKKβ (Fig. 4D, S14). The ratio of this slowly migrating A20 species to the more rapidly migrating form was markedly decreased in blood leukocytes from healthy donors heterozygous for the Denisovan T108A;I207L variants, and further decreased in a healthy T108A;I207L homozygous donor, compared to healthy controls without these variants (Fig. 4E). The ratio of phosphorylated-S381 to fast-migrating A20 was also decreased (Extended Data Fig. 14F). TNF-α stimulated leukocytes from variant carriers also had heightened expression of NF-κB-induced transcripts of *ICAM1*, *CXCL2, TNF*, and *TNFAIP3* itself (Extended Data Fig. 15), and a trend towards reduced IκBα protein levels (Extended Data Fig. 14H).

When A20 was measured for inhibition of an NF-κB luciferase reporter in unstimulated or TNFα-stimulated mouse insulinoma cells, inhibition was diminished in a graded fashion by each OTU domain variant in the same order that these diminished S381 phosphorylation (Fig. 4F; Extended Data Fig. 13C). The deleterious C243Y variant was the most compromised inhibitor of the series, almost as compromised as A20 with S381 substituted to non-phosphorylatable alanine (S381A). Combining the intermediate I325N mutation with a S381A mutation *in cis* did not cause a further decrease in A20 inhibition, consistent with I325N having its effect upon phosphorylation. That conclusion was reinforced by combining I325N *in cis* with a substitution of S381 to the phosphoserine mimetic glutamate (S381E), which rescued the lost activity caused by I325N (Extended Data Fig. 13D, E). Testing the two Denisovan variants individually, T108A had no measurable effect whereas I207L diminished A20 inhibitory activity comparably to the two variants combined (Extended Data Fig. 13F), supporting I207L as the functional variant within the introgressed Denisovan haplotype.

To explore the structural and biochemical consequences of variants on the posterior surface of the OTU domain, we focused on the intermediate I325N allele. Crystallographic structures of A20 OTU domains with wild-type I325 or mutant N325 revealed no differences in features with known functions, including the catalytic triad and ubiquitin-binding surface (Fig. 1B; Extended Data Fig. 16A-C). I325N also did not alter the conserved posterior surface of the OTU domain^40^, including the β3-β4 loop containing C243 (Extended Data Fig. 16G), although there were subtle shifts in the stem of the β7-β8 loop, which contains conserved surface residues T321, T322 and L324 (Extended Data Fig. 16D-G). The β7-β8 loop itself is disordered in all available OTU structures but, like the disordered loops in the unliganded S1 ubiquitin-binding site, it may undergo conformational changes upon binding a cognate partner that are hindered by the I325N variant. Wild-type and variant OTU domains exhibited similar thermal denaturation profiles (Extended Data Fig. 17A-C), and the corresponding full-length proteins were comparably stable in cycloheximide-treated mammalian cells (Extended Data Fig. 17D). I325N did not decrease DUB activity of bacterially expressed OTU domain against K48-polyubiquitin *in vitro* (Extended Data Fig. 17E-G), but I325N diminished K63-polyubiquitin DUB activity and K48-ubiquitin ligase activity when full length A20 was expressed in human cells (Extended Data Fig. 17H, I) consistent with these activities requiring S381 phosphorylation^18,19^.

## Shift from beneficial to detrimental loss of microbial tolerance

We next explored the possibility that partial loss of A20 phosphorylation and subclinical loss of microbial tolerance might become detrimental in particular environmental or genetic settings. In an experimental model for septic shock, I325N homozygous mice had increased IL-6 production and high mortality following injection of a dose of bacterial LPS that was not lethal in wild-type controls (Fig. 5A, B). In the rodent counterpart of smallpox, infection with the orthopoxvirus ectromelia virus was tolerated and controlled by wild-type mice, yet resulted in high mortality and higher viral titres in I325N homozygotes (Fig. 5C, D and Extended Data Fig. 18). In bone marrow chimeric mice with a higher frequency of pancreatic islet-reactive CD4^+^ T cells, autoreactive T cells escaped deletion and precipitated diabetes when half the hematopoietic cells were homozygous for I325N (Extended Data Fig. 19A-C). The rogue islet-reactive T cells were nevertheless derived equally from A20 wild-type or I325N precursors, whereas I325N acted cell autonomously to increase MHC II and the T cell costimulator CD86 on B cells and dendritic cells and to increase frequencies of germinal centre B cells (Extended Data Fig. 19D-F).

**Fig. 5.**
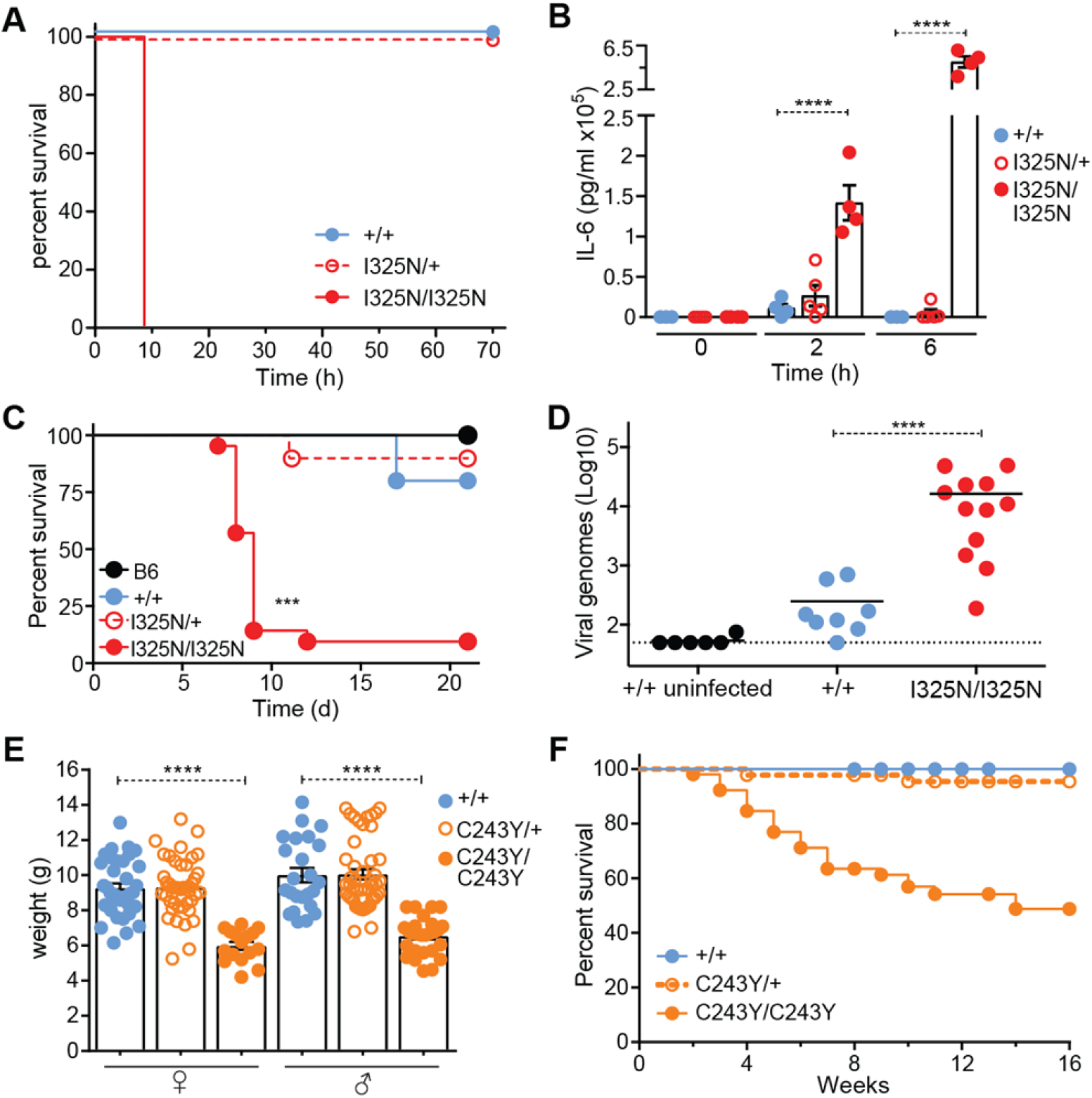
Shift from beneficial to detrimental effects in response to environmental or genetic stressors. Kaplan-Meier survival curves **(A)** and serum IL-6 concentration **(B)** in mice of the indicated genotypes given 50 µg LPS by intraperitoneal injection at time=0. *N*=16-18 for each curve in (A). **(C)** Kaplan-Meier survival curves for sex-matched littermates of the indicated *Tnfaip3* genotypes and wild-type controls (B6) infected with Ectromelia virus. **(D)** Day 8 serum viral load assessed from mice in (C). (**E**) Weight of 3-week-old *Tnfaip3*^C243Y/C243Y^ mice and heterozygous or wild-type littermates. **(F)** Survival of C243Y homozygous mice (*n*=52) compared to heterozygous (*n*=43) or wild-type littermates (*n*=41). Statistical analysis by Students *t*-test and one-way ANOVA: \*\*\**P* < 0.001, \*\*\*\**P* < 0.0001.

While specific stressors were required to reveal detrimental effects of the intermediate I325N variant, wasting disease occurred spontaneously in young C57BL/6 mice homozygous for the severe C243Y variant. C243Y homozygotes were born at the expected Mendelian ratio but compared to wild-type littermates failed to thrive, exhibiting a large reduction in body weight and early lethality (Fig. 5E, F; Extended Data Fig. 20). Surviving C243Y homozygotes showed intestinal pathology including loss of goblet cells responsible for sustaining the mucin barrier between epithelium and microbes of the intestinal lumen, along with an increased frequency of circulating CD44^hi^ activated/memory T cells (Extended Data Fig. 20). C243Y homozygous mice thus resemble A20-null mice^5,11^, paralleling the clinical similarity between C243Y missense and A20 loss-of-function variants in heterozygous humans^12^.

## Discussion

Our findings reveal genetic and biochemical mechanisms for adaptively increasing immunity, with trade-offs against microbial tolerance and anabolic metabolism that either remain clinically silent or become detrimental in specific contexts. Three different missense alleles in the N-terminal A20 OTU domain act, to different degrees, by diminishing A20 phosphorylation and tuning down A20’s immune inhibitory activity. A rare human variant, C243Y, diminishes phosphorylation almost entirely and shifts the balance away from microbial tolerance to the extremes of immunity, resulting in severe inflammatory disease in both mice and humans. This outcome is comparable with other human variants that eliminate or truncate A20 protein from one *TNFAIP3* allele or engineered mouse variants that eliminate A20 from both alleles^11,12^. By contrast, I325N and I207L variants decrease A20 phosphorylation more modestly, without precipitating spontaneous inflammatory disease in the mice or humans who harbor them. I325N in the mouse confers dramatically increased resistance to an otherwise lethal dose of a Coxsackievirus. A similar resistance to microbial pathogens may explain the beneficial effect of Denisovan I207L, as evidenced by its high frequency in modern human populations east of the Wallace Line, who likely encountered new infectious agents as they moved into environments with unique fauna and flora.

Previous studies in mice have conditionally deleted A20 in specific tissues, causing severe inflammatory disease, or directly disabled individual enzymatic functions of A20 in all tissues resulting in surprisingly little inflammation^5,6,11,15,18,20^. By contrast, OTU domain variants tuning phosphorylation regulate multiple ubiquitin-editing functions of A20^18,19^, and as shown here have greater effects on microbial tolerance and resistance *in vivo* than variants affecting individual ubiquitin-editing activities. The C243Y and I325N variants involve buried residues in two separate surface loops, β3-β4 and β7-β8 respectively (Extended Data Fig. 16), on the highly conserved posterior surface of the OTU domain^40^. Structural studies of full-length A20 protein may illuminate how this surface relates to the C-terminal domain carrying S381. Because the conserved surface bounded by C243Y and I325N is large, it potentially offers many opportunities for substitution of buried or surface residues to tune immunity and tolerance.

Our study of the intermediate loss-of-function A20 variant, I325N, provides two examples of genetic trade-offs. The first was with glucose metabolism, where *Tnfaip3*^I325N^ had the surprising effect within pancreatic islet beta cells of decreasing insulin secretion while increasing inflammation. This is reminiscent of findings in *Drosophila* that insulin production or action decreases during infection, diminishing body glycogen supplies through increased FOXO activity, a transcriptional inducer of starvation responses^37^. Resting insulin levels are a good prognostic marker in human sepsis, and exogenous insulin treatment improves outcome^41^. Subtly decreased A20 activity may enhance immune demands for energy and, separately, help meet these energy demands by lowering insulin-induced anabolic growth, both contributing to cachexia associated with chronic infection and inflammation.

The second example of a trade-off is the experimental demonstration that a beneficial trait in one context can be deleterious in another. *Tnfaip3^I325N^* conferred increased resistance to Coxsackievirus, an important human enterovirus, but increased mortality to ectromelia virus, a relative of variola and myxoma viruses. Higher ectromelia virus mortality in *Tnfaip3^I325N^* mice is reminiscent of the high mortality in indigenous populations of the Americas and Oceania exposed to variola virus relative to Europeans^8^. The findings here suggest that the devastation wrought by these and other microbes reflects selection during earlier environmental conditions for lower microbial tolerance and a stronger immune response, with A20 representing one critical determinant of this outcome.

## Acknowledgments

This manuscript is dedicated to the memory of Susan Watson, who led the isolation of the I325N mouse. We wish to thank the National Computational Infrastructure, Xenon systems, Nvidia, and the Multi-modal Australian ScienceS Imaging and Visualisation Environment (MASSIVE) (www.massive.org.au) for compute resources. We thank Oliver Venn, Iain Mathieson, Bastien Llamas, Yassine Souilmi, Raymond Tobler and Alan Cooper for stimulating discussions. CNJ thanks Jason Cyster for mentorship. We thank Wendy Sandoval for assistance with A20 phosphorylation site mass spectrometry, and Kim Newton and Vishva Dixit for reagents. This study makes use of data generated by the Telethon Kids Institute. A full list of the investigators who contributed to the generation of the data is available from http://bioinformatics.childhealthresearch.org.au/AGHS.

## Author contributions

Human family recruitment and genome sequence analysis: PG, JBZ, MEC, PB-A, JT, MW, OMS, AR, MJC, VG, MED, CCG. Tracing Denisovan haplotype in human populations by OMS, PG, AR, MPC, CCG; peripheral blood analyses by NWZ, AR, CCG, STG. I325N mutation: identification by SRW, CNJ, LLL, TW, CR, BW, CCG; immune cell analysis by KH, SRD, YJ, NWZ, MY, EB, AE, MHS, GPN, FS, DEZ, AA, IEW, STG, CCG; Coxsackievirus studies by NWZ, CL, JW, CK, STG; Ectromelia studies by GC, GK; body weight, islet, LPS and inflammation studies by NWZ, SNW, EKM, JEV, STG; protease activity and structural studies by KH, NWZ, DBL, DC, WP, IEW, BEC, PDM, CJJ, BTP, AMB, STG, CCG. C243Y mouse strain production and analysis by NWZ, RB, DZ, SNW, DC, JW, STG. A20 phosphorylation and NF-κB luciferase analysis by NWZ, DC, DWA, WP, IEW, CCG, STG.

## Data and materials availability

All data is available in the main text or the supplementary materials.

## Materials and Methods

### Human subjects

Patients and healthy family members were recruited via the Clinical Immunogenomics Research Consortium Australia and The Children’s Hospital at Westmead, providing written informed consent under protocols approved by relevant human research ethics boards.

### Genome sequencing

Parent-proband trio genomes were sequenced on the Illumina HiSeq X platform using DNA isolated from whole blood. Libraries were generated using either Illumina TruSeq PCR-free or TrueSeq Nano.

Raw reads were aligned to the hs37d5 reference using BWA-MEM v0.7.10-r789 (arXiv:1303.3997 [q-bio.GN]), and sorted and duplicate-marked with Novosort v1.03.01 (Novocraft Technologies). The GATK suite v3.3-0-g37228af ^43^ was used for local indel realignment and base quality score recalibration. gVCFs were generated with GATK HaplotypeCaller, joint-called as trios using GATK GenotypeGVCFs, and variants recalibrated using GATK Variant Quality Score Recalibrator (VQSR). Finally, VCF files were annotated with Variant Effect Predictor (VEP) v76^44^ using the LoFTEE^45^ and dbNSFP plugins.

Variants were packaged into databases using GEMINI (v0.18.3)^46^ and imported into Seave^47^. All reported variants were manually inspected using IGV^48^ to verify their authenticity and validated by Sanger Sequencing.

### Population genetic analysis

*TNFAIP3* genotypes were extracted from 279 publicly-available genomes from the Simons Genome Diversity Project samples^22^, or exome data from 72 Martu indigenous Australians^32^. In addition, a collection of 514 individuals from across mainland and Island Southeast Asia, Papua New Guinea and Oceania was genotyped at 567,096 variants using the Affymetrix Axiom Genome-Wide Human Array, with 538,139 autosomal variants with <5% missing data kept for further analyses^30^. Weir and Cockerham’s estimated F_ST_ values were calculated using VCFtools v0.1.14, and principal component analysis was performed using PLINK v1.9. Genotypes were phased and imputed using the Michigan Imputation Server v1.0.3 (http://imputationserver.sph.umich.edu), and compared to high-coverage Denisovan^27^ and Altai Neanderthal^28^ *TNFAIP3* haplotypes. Phred-scaled CADD scores^49^ were calculated for all PASS variants across the extended Denisovan haplotype with gnomAD allele frequency <0.01.

### PBMC isolation and manipulation

Human PBMCs were prepared using a Ficoll-Hypaque gradient and live count taken using 0.1 % trypan blue. Cells were resuspended at 10^6^ cells/ml in sterile freezing medium (RPMI-1640, 10% DMSO, 50% FCS) in 1.5 ml Sarstedt cyrovials and stored in liquid nitrogen. Human PBMCs were thawed in a 37°C water bath and added drop wise to pre-warmed medium (RPMI-1640, 10% FCS, 100 U/ml P/S, 1% HEPES, 2 mM L-Glutamine, 2mM EDTA) and resuspended for culture at 37°C + 5% CO_2_ for 1 h prior to stimulation with recombinant human TNFα (R&D Systems). Following stimulation, cells were pelleted and processed for immunoblotting or RT-qPCR, as described below.

### Mice

The *Tnfaip3 Lasvegas* strain (*Tnfaip3*^I325N^) were generated by *N*-ethyl-*N*-nitrosourea (ENU) mutagenesis of C57BL/6 mice, and propagated by backcrossing to C57BL/6. The strain was maintained as heterozygous breeding pairs so that WT littermates could be used for controls. To generate *Tnfaip3^C243Y^* mice, a guide RNA with the sequence 5′- GGGATATCTGTAACACTCC-3′ was microinjected into C57BL/6 zygotes in combination with Cas9 mRNA. Founder mice carrying the C243Y substitution were then crossed to C57BL/6, and heterozygous offspring intercrossed to generate homozygous, heterozygous and wild-type littermates for experimentation. Mouse lines were housed at the Australian Phenomics Facility (Australian National University, Canberra, Australia), or at the Australian BioResources Centre (ABR) (Moss Vale, NSW, Australia). *Tnfaip3*^C103A 18^, 3A9 TCR transgenic^50^ and insHEL transgenic^51^ mice have been described. Mice were genotyped for transgenes and mutations by PCR and used 7–28 (typically 10-16) weeks after birth. For TCR^3A9^ transgenic mouse experiments, mice were hemizygous for the 3A9 or insHEL transgenes on the B10.BR and the B10.BR.SJL-*Ptprc^a^* (CD45.1) backgrounds, and the *Tnfaip3*^I325N^ mutation was backcrossed to B10.BR. To make chimeras, congenic B6.SJL- *Ptprc^a^* or B10.BR.SJL-*Ptprc^a^* (CD45.1) mice were irradiated with two doses of 4.5 Gy 4 hrs apart, and injected IV with mixtures of 1.8×10^6^ bone marrow cells from B6 or B10.BR SJL-*Ptprc^a^* mice and 1.8×10^6^ bone marrow cells from B6 (CD45.2) mutant or control mice, and analyzed 8-14 weeks later. Animal studies were approved by the Garvan/St Vincent’s or the Australian National University Animal Ethics Committees. All procedures performed complied with the Australian code of Practice for Care and Use of Animals for Scientific Purposes.

### Coxsackievirus infection model and virus quantification

Mice were intraperitoneally injected with normal saline (control) or 20 plaque forming units (PFU) in 200 µl saline of the CVB4 E2 lab strain kindly provided by Prof. Malin Flodström Tullberg (Karolinska Institutet, Solna, Sweden) grown in HeLa cells. Mice were monitored daily and euthanized if they displayed gross signs of illness (e.g., ruffling, hunching). The pancreas was harvested at indicated times for RT-qPCR and histopathology analysis and viral titre determination. Serum was also collected by cardiac puncture for measurement of IL-6 by ELISA (BD; OptEIA^TM^ Set Mouse IL-6), as per manufacturer’s instructions. Non-fasting blood glucose levels were measured at least twice-weekly using Freestyle Lite Blood Glucose Test Strips (Abbott, Australia). Plaque assays were performed to determine viral titres within the pancreas following infection. HeLa cells (0.6 × 10^6^/well) were seeded in 2 ml complete medium (RPMI with 10% FCS, 2mM L-Glutamine, 100 U/ml Penicillin, 100 µg/ml Streptomycin and 100 µg/ml Normocon) in 6 well plates and incubated over night at 37°C. Infections samples were collected in RPMI, homogenized in a Dounce tissue grinder and passed through a 22 µm filter, before preparing tenfold serial dilutions. HeLa cells, at 90% confluency, were washed with 1X PBS and 400 µl of infectious homogenate added to each well and incubated for 60 min at 37°C under gentle rocking. Infectious media was removed and 3 ml of agar mix (2x 1.8% agar, 2x MEM containing 10% FCS) was added to each well before incubating at 37°C for 3 days. Cells were then fixed with Carnoys reagent for 60 min and subsequently stained with 0.5% Crystal violet for 60 s. Wells were extensively washed with H_2_O and plaques counted.

### Ectromelia virus model

Mice were inoculated subcutaneously with 10^3^ PFU Ectromelia virus (Moscow strain; ATCC #VR-1374) in the flank of the left hind limb between the regio tarsi and regio pedis under avertin anesthesia. Clinical scores were recorded daily, and animals weighed on days 0, 5 and every 2 days thereafter until day 21, and bled on days −1 and 8 to measure viral load. Animals with a significant clinical score or 25% decrease of starting weight were euthanized and considered dead the following day. Viral load was measured in blood by qPCR for *ECTV-Mos-156* viral genomes and in organs by viral plaque assay as log_10_ PFU/gram tissue^52^.

### Macrophage cultures

Bone marrow from femurs and tibiae was cultured 7 days in complete RPMI1640 medium with 10% fetal bovine serum, 100U/ml penicillin, 100µg/ml streptomycin, 2mM L-glutamine (Life Technologies) and 50ng/ml recombinant human Macrophage colony-stimulating factor (Peprotech). Cells were placed into 6-well plates for stimulation with 10ng/ml Salmonella Minnesota R595 (Re) ultra-pure LPS (List Biological) for different time-points. RNA was extracted by TRIzol (Life Technologies), reverse transcribed to cDNA with SuperScriptII (Life Technologies), *Cxcl1*and *Cxcl11* levels were measured using the Taqman assay (Applied Biosystems). RNA levels relative to *Ef1a* were calculated with the delta Ct values. Conditionally *Hoxb8*-immortalized bone-marrow progenitor cells were generated as described^53^.

### LPS sepsis model

A low dose of 50µg LPS was administered by intraperitoneal injection. Mice were monitored every 1 h for 10 h and sacrificed if ethical end-point reached. Monitoring was conducted using body conditioning score and Grimace Scale (NC3Rs [National Centre for the Replacement Refinement & Reduction of Animals in Research) approved by the Garvan/St Vincent’s Animal Ethics Committee. Monitoring continued twice daily for 7 days in surviving mice, were weights and blood glucose levels were also measured. Serum was collected by tail-tipping before LPS injection and 2 and 4 hours following injection for IL6 determination by ELISA (BD; OptEIA^TM^ Set Mouse IL-6), as per manufactures instructions.

### Immunohistochemistry and beta cell area determination

Tissues were fixed in 10 % neutral buffered formalin (Sigma-Aldrich), paraffin embedded and parallel sections (5 µm) prepared. Sections were stained with hematoxylin and eosin (H&E; Sigma-Aldrich) and for pancreatic tissue parallel sections stained for insulin (purified rabbit anti-mouse insulin polyclonal antibody; 4590; Cell Signaling Technology). Visualization of bound anti-insulin antibody was achieved using HRP-labelled polymer-conjugated goat anti-rabbit IgG (Dako EnVision+ System), followed by counterstain with hematoxylin. For pancreatic beta cell mass determination consecutive pancreatic serial sections at 200 µm intervals were stained for insulin and beta cell area quantified from total area taken by insulin positive cells compared to non-positive tissue (ImageJ). Beta cell mass (mg) was calculated by multiplying relative insulin-positive area by the mass of the isolated pancreas before fixation. Images were captured using a Leica DM 4000 or Leica DM 6000 Power Mosaic microscope (Leica Microsystems).

### Metabolic studies

Blood glucose levels (BGL) were determined using a FreeStyle Lite® glucometer and blood glucose test strips (Abbott Diabetes Care) via tail tipping. Measurements were taken from 8 and 12 week old male or female non-fasted mice. Intraperitoneal glucose tolerance tests (IP-GTT) were conducted following an overnight fast (16 h) with access to water. The following day mice were weighed and fasting blood glucose measurements taken. Subsequently, mice were injected (IP) with 20% dextrose w/v (Sigma Aldrich) to a final concentration of 2 g glucose per kg body weight (2g/kg). Blood glucose levels were measured from the tail vein at 15, 30, 60 and 120 min post-glucose administration. Intravenous (IV) GTT was conducted in a similar manner; however, glucose (1 g/kg) was administered intravenously into the tail vein and blood glucose measurements taken at 0, 5, 10, 15, 20, 30, and 60 min post-injection. During the IV-GTT blood samples were also taken for the determination of insulin content via ELISA, conducted as per manufactures instructions (Cayman Chemical). Glucose-stimulated insulin secretion assay (GSIS) was performed for islets *ex vivo* as descried^54^.

### Islet isolation, transplantation, and *in vitro* studies

Islets were isolated as previously described^55^, and counted for islet transplantation or *in vitro* experiments using a Leica MZ9.5 stereomicroscope. Islets were transplanted under the kidney capsule of diabetic B6 littermates as described^56^. Diabetes was induced by IP injection of 180 mg/kg streptozotocin (Sigma-Aldrich) dissolved in 0.1 M citrate buffer (pH 4.2) at a concentration of 20 mg/ml. Diabetes was determined as [blood glucose] ≥16 mM on two consecutive days measured by FreeStyle Lite® glucometer and Abbott Diabetes Care test strips following tail tipping. Islet grafts were retrieved from recipients at indicated time points post-transplantation for analysis of islet morphology, function, or degree of lymphocytic infiltration by histology or gene expression by RT-qPCR. Gene expression in islet grafts was calculated using the average WT ΔCt value. Islets to be used for *in vitro* studies were cultured overnight in islet overnight culture media (RPMI-1640, 20% FCS, 100 U/ml P/S, 2 mM L-Glutamine) at 37°C + 5% CO_2_.

### Flow cytometry

Flow cytometric staining was performed as described in^57,58^. Antibodies against the following surface antigens were: CD4 (RM4-5), CD8 (53-6.7), CD21 (7G6), CD23 (B3B4), CD25 (PC61.5), CD44 (IM7), CD69 (H1.2F3), CD93 (AA4.1), CD45.2 (104), B220 (RA3-6B2), IgM (II/41), IgD (11-26). Splenocytes were labeled with CFSE (Invitrogen) and cultured in RPMI1640 supplemented with 10% FCS, 2 mM L-glutamine, 55 µM 2- mercaptoethanol, and penicillin/streptomycin. B cells were stimulated with: F(ab’)_2_ goat anti-mouse IgM (10 µg/ml, Jackson Immunoresearch Laboratories), LPS from *E. coli* 055:B5 (10, 1, 0.1 µg/ml, SIGMA), anti-CD40 (10 µg/ml FGK4.5, BioXCell). When analysing 3A9 TCR transgenic cells single cell suspensions from thymus, spleen or pancreatic lymph node were incubated for 30 min at 4° C in culture supernatant from the 1G12 hybridoma (specific for TCR^3A9^). Then, cells were pelleted by centrifugation and incubated for 30 min at 4° C in FACS buffer containing assortments of fluorochrome- or biotin-conjugated monoclonal antibodies against cell surface proteins. After washing in FACS buffer, cells were fixed and permeabilised using the eBioscience Foxp3 staining buffers, then incubated with antibodies specific for intracellular proteins and Brilliant Violet 605-streptavidin conjugate (BioLegend) to detect biotin-conjugated antibodies. Antibodies were purchased from BD, eBioscience or BioLegend. Data were acquired with LSR II flow cytometers (BD) and analyzed using FlowJo software (Tree Star).

### CyTOF

Unstimulated spleen cells from four wild-type and four I325N-homozygous mutant mice were individually labeled with mass-barcodes, mixed, permeabilized and stained with mass-labeled antibodies to a panel of cell surface markers and intracellular proteins including IκBα, and analyzed by CYTOF mass spectrometry^59,60^. Spanning-tree Progression Analysis of Density-normalized Events (SPADE)^61^ analysis was used to resolve leukocyte lineages and subsets, and the relative intensity of IκBα in each subset compared between mutant and wild-type cell counterparts.

### Real Time quantitative PCR (RT-qPCR)

Mouse islets were isolated and placed into 12-well non-tissue culture-treated plates (150- 200 islets/well; Fisher Scientific). Following an overnight culture islets were treated with 200 U/ml recombinant human (h) TNFα (R&D Systems) for 1, 4, or 24 h. Total RNA was extracted using the RNeasy Plus Mini Kit (Qiagen) and reverse transcribed using Quantitect Reverse Transcription Kit (Qiagen). Primers were designed using Primer3 software^62^ with sequences obtained from GenBank and synthesized by Sigma Aldrich (Table 3, 4). PCR reactions were performed on the LightCycler^®^480 Real Time PCR System (Roche) using the FastStart SYBR Green Master Mix (Roche). Cyclophilin (CPH) was used as the housekeeping gene and data analyzed using the 2^ΔΔCT^ method. Initial denaturation was performed at 95° C for 10 sec, followed by a three-step cycle consisting of 95° C for 15 sec (4.8° C/s, denaturation), 63° C for 30 sec (2.5° C/sec, annealing), and 72° C for 30 sec (4.8° C/s, elongation). A melt-curve was performed after finalization of 45 cycles at 95° C for 2 min, 40° C for 3 min and gradual increase to 95° C with 25 acquisitions/° C. Expression differences were visualized using GraphPad or BAR Heatmapper plus tool.

### Immunoblot analysis and immunoprecipitation

Primary islets were lysed in islet lysis buffer (50 mM Tris-HCL pH7.5, 1% Triton X, 0.27 M sucrose, 1 mM EDTA, 0.1 mM EGTA, 1 mM Na_3_VO_4_, 50 mM NaF, 5 mM Na4P2O7, 0.1% β-mercaptoethanol; supplemented with EDTA-free protease inhibitor [Roche]), βTC_3_ cells were lysed with radioimmunoprecipitation (RIPA) buffer with SDS, following relevant treatment with or without 200 U/ml of recombinant human (h) or mouse (m) TNFα (R&D Systems). Protein concentration was measured using the Bradford assay (Bio-Rad) and total protein (20-25 µg) resolved on a 7 - 10% SDS PAGE gel and then transferred to a nitrocellulose membrane, Immobilon-P® (Merck Millipore). Membranes were incubated with anti-IκBα (9242), anti-phospho-IκBα (9256), anti-JNK (9252), anti-phospho-JNK (9255), anti-IκKα (2682), anti-phospho-IκKα/β (16A6; 2697), anti-NIK (4994), anti-NF-κB2 p100/p52 (4882), anti-RelB (C1E4, 4922) (Cell Signaling Technology); anti-beta-actin (AC15) (Sigma-Aldrich); or anti-S381-A20 (a kind gift by Professor Derek W. Abbott^19^) followed by horseradish peroxidase (HRP)-labeled secondary antibody goat-anti-mouse IgG Fc (Pierce Antibodies) or donkey-anti-rabbit IgG (GE Life Sciences). HRP conjugates bound to antigen were detected and visualized by using an ECL detection kit (GE Life Sciences).

Immunoprecipitation was conducted in thymocytes, which were lysed at 4°C in immunoprecipitation buffer (0.025M Tris, 0.15M NaCl, 0.001M EDTA, 1% NP-40, 1% Triton × 100, 5% glycerol) containing Complete® protease inhibitor and Phosphostop® phosphatase inhibitors (Roche). Immunoprecipitation was then conducted by first preclearing lysates with protein A/G-Sephrose (Thermo Fisher Scientific) for 1 h and then incubated with anti-TNFR1 (ab7365) cross-linked beads or anti-A20 (59A426) antibody (Abcam) for 2 h at 4°C. Following incubation with only antibody 25 μl of protein A/G beads were added and incubated at 4° C on a roller overnight. Beads were washed 4 × with lysis buffer and then eluted using low pH amide buffer for cross-linked beads or 30 μl of Laemmli reducing gel-loading sample buffer for non-cross linked beads. Samples were vortexed, heated to 100° C for 5 min, cooled on ice for 10 min, and then loaded onto a 8 or 10% agarose gel for immunoblotting for anti-A20 (56305/D13H3), anti-IκBα (9242), anti-IκKβ (2684), anti-JNK (9252), anti-TAK1 (52065), anti-phospho-IκBα (9256), anti-phospho-JNK (9255), anti-phospho-TAK1 (4536/90C7), anti-TNFR1 (3736C25C1) (Cell Signaling Technology), anti-RIP1 (H-207) (Santa Cruz), anti-RIP1 (610458) (BD bioscience), anti-TAK1 (491840) (R&D systems), or anti-beta-actin (AC15) (Sigma-Aldrich). For B cell stimulation, splenic B cells purified by MACS with CD43 depletion were stimulated with anti-IgM F(ab’)2 or LPS for the indicated times. Cells were lysed with TNE buffer (1% Nonidet P-40, 20 mM Tris-HCl, pH 8.0, 150 mM NaCl, 0.1 mM sodium orthovanadate, and complete protease inhibitor [Roche]). The following antibodies were used: anti-IκBα (9242), anti-TNFAIP3 (5630/D13H3) (Cell Signaling Technology), anti-A20 (A-12), anti-ubiquitin (P4D1) (Santa Cruz) and anti-beta-actin (AC-15) (Sigma-Aldrich).

### *In vitro* reporter transfection studies

Reporter assays were carried out as described previously^63,64^. NF-κB activity experiments were conducted using βTC_3_ cells transfected with 0.3 μg of the NF-κB.Luc reporter (Promega) and 0.25 μg CMV.β-galactosidase (a kind gift from Beth Israel Harvard Medical School, Boston, MA). pcDNA vectors encoding human WT or variant A20 constructs or the empty pcDNA3.1 reporter were then added (0.3 μg), and each well topped with 0.15 pcDNA3.1 to make 1 μg total DNA. Transfection was conducted using Lipofectamine 2000 (Invitrogen). Following transfection cells were stimulated with 200 U/ml of recombinant human (h) TNFα (R&D Systems). Luciferase activity was assayed in cell lysates harvested 8 h post-stimulation, using a luciferase assay kit (Promega). Results were normalized to β-galactosidase activity (Galactostar) to give relative luciferase activity. Expression plasmids and reporters were obtained and maintained as described previously^63,64^.

### Ubiquitination assays

Wildtype A20, A20^I325N^ or A20^C103A^ OTU domain protein (1 µg) purified from *E. coli* was added to 2 µg of K48-ubiquitin chains of mixed chain length (Ub_2_-Ub_7_) or to purified tetra-ubiquitin (Ub_4_) (#UC-230, UC-210, Boston Biochem). A20 OTU domains added to mixed-length K48-ubiquitin chains were incubated in 100 µl of deubiquitin buffer (50 mM HEPES pH 8.0, 0.01% Brij-35, 3mM DTT) at 37° C with agitation at 400 rpm for 60 or 150 min in a benchtop incubator shaker. A20 OTU domains added to K48-tetra-ubiquitin chains were incubated in 100 µl of deubiquitin buffer (25 mM HEPES pH 8.0, 5 mM DTT, 5 mM MgCl_2_). At the indicated times, 20 µl of reaction mixture was collected and the enzymatic reaction stopped by addition of SDS sample buffer. Recombinant Flag-tagged full length A20^+/+^ or A20^I325N^, was expressed in HEK-293T cells and purified as described previously^14,17^. Full-length A20 deubiquitination reactions were performed using 100ng recombinant A20, 500 ng of the indicated ubiquitin chain and DUB reaction buffer with or without phosphatase inhibitor cocktail and were incubated for the indicated times at 37° C with agitation at 1,000 rpm. Following incubation samples were placed on ice and 20 µl collected and added to SDS sample buffer to stop the reaction. Full-length ubiquitin ligase assay was performed as previously described^14^. Samples were subjected to 1D SDS-PAGE and immunoblotted for ubiquitin (clone P4D1; Santa Cruz or Cell Signaling Technology) or A20 (clone 59A426; Abcam or A-12; Santa Cruz), as described above.

### OTU protein preparation and crystallisation

The N-terminal OTU domains of human and mouse A20 (human WT and I325N, residues 1-366; mouse WT and C103A, 1-360) were cloned into vector pGEX-6P-1 (GE Healthcare), facilitating bacterial expression as a GST-fusion. Sequences were confirmed by Sanger sequencing. Expression was performed in *E. coli* strain BL21 (DE3) Gold, where cells were induced at an OD600 of 0.5 with 0.2 mM IPTG, followed by incubation at 20° C overnight in LB medium. Cells were harvested by centrifugation, lysed by three cycles of freeze-thaw and one cycle of pressure shock (human WT and I325N) or by sonication alone (mouse WT and C103A). The GST-fusions was captured from the cleared lysate by passage over GSH-Sepharose CL4B resin (GE). For purification of human WT and I325N, the resin was washed (50 mM Tris (pH 8.8) 200 mM NaCl, 5 mM DTT, 1 mM EDTA), then the A20 component released by overnight incubation (4° C) with PreScission protease. Eluted A20 was stabilized by incubation with iodoacetamide (Sigma; 30 mM, 30 min at RT). The reaction was terminated by addition of an equivalent amount of β-mercaptoethanol. For purification of mouse WT and C103A protein, purified GST-OTU fusion protein was eluted (50 mM Tris-HCl [pH 8.0], 200 mM NaCl, 5 mM DTT, 10 mM glutathione) from the column and the A20 OTU component was released by overnight incubation (4° C) with PreScission protease. The cleavage products were subject to anion-exchange chromatography (HiTrap Q FF; GE) where the OTU domain eluted as a single peak between 5 and 500 mM NaCl, 25 mM Tris-HCl pH 7.5. All proteins were then concentrated and subject to gel filtration chromatography (ÄKTA S200 26/60, buffers as described above).

Crystals of both the A20^I325N^ variant (long triangular rods) and wild-type A20 (long rods) OTU domains, grown under the same conditions; equal volumes of protein (2.7 mg/mL) and well solution (50 mM CaCl_2_, 100 mM MES (pH 6.0), 5% PEG1500) were combined in a hanging drop setup. Crystals grew over several weeks at RT. Partial cryoprotection was achieved by briefly (1-5 seconds) swimming crystals in a solution comprising equal volumes of mother liquor and well solution doped with glycerol (25% v/v final) prior to being plunge vitrified in liquid N_2_. For the mouse wild-type A20, crystallization was achieved by vapor-diffusion in hanging-drops at a protein concentration of 8 mg/mL in 1.8 - 2.4 M NaCl, 0.1 M MES [pH 6-6.7]. Crystals were soaked in mother liquor containing 30% ethylene glycol for one minute and immediately vitrified in a nitrogen cryo-stream.

### Crystallographic data reduction and model refinement

Diffraction data using light of wavelength 0.9537 Å was collected at 100 K at beamline MX2 at the Australian Synchrotron. Data were indexed and integrated with MOSFLM^65^. The spacegroups were scrutinized with POINTLESS, and the data scaled with AIMLESS^66,67^, or SCALA^68^ accessed via the CCP4i software interface^69^. Although grown under essentially the same conditions, the human WT and I325N mutant A20 proteins crystallized in different space groups. These data were highly anisotropic, resulting in poor completeness, low multiplicity, and noisy electron density maps. In the case of the mouse OTU crystal, there was significant thermal diffuse scattering. See Table 5 for data reduction and refinement statistics.

Structures were solved by molecular replacement using PHASER^70^. The search model was the A-chain of PDB entry 3DKB, but stripped of surface loops that displayed conformational variability in other PDB entries (2VFJ and 3ZJD). In the case of the human I325N data the structure was originally solved in the space-group P3_1_2 with 4 molecules in the asymmetric unit. However, crystal packing and residual unaccounted-for electron density suggested more molecules might be present. The structure was subsequently solved in the lower symmetry P3_1_ space group, with 6 molecules in the asymmetric unit, sensible packing and no unaccounted-for density. The human wild-type A20 structure, solved as a control for the iodoactamide alkylation, also has 6 molecules (three dimers) in a different asymmetric unit and space group. The mouse OTU structure was indexed and refined in the P3_2_ space group, with a dimer in the asymmetric unit. Restrained B-factor refinement, using local non-crystallographic symmetry (NCS) restraints, was performed with REFMAC5^71^. For the mouse A20 OTU structure TLS and restrained refinement was carried out using phenix.refine^72^. Between rounds of refinement, electron density maps and composite OMIT maps^73^ and their fit to the model were examined using COOT^74^. Amino acid side chains were added/subtracted if suggested by difference map electron density. The active-site cysteine (C103) was clearly identified through inspection of mFo-Dfc difference maps as the only cysteine residue alkylated by the iodoacetamide treatment in both human structures (PDB residue descriptor YCM). All 6 molecules refined in each structure are highly similar in fold, both to themselves, and compared with each other. All molecules form dimers with neighboring molecules, as observed in other crystal structures. Structure validation was performed using the MOLPROBITY web server^75^. The final human I325N, human WT, and mouse WT structures contain Ramachandran favored/outlier components of 88.64/0.00%, 88.13/2.87 %, and 82.13/7.87 %, respectively.

### Measurement of OTU domain thermal stability

Purified A20 OTU domains (0.4 mg.mL^-1^) were exchanged into 100 mM NaCl, 5 mM dl-dithiothreitol, 20 mM Na_2_HPO_4_ (pH 7.5). Circular dichroism data were collected with a Chirascan circular dichroism spectrometer (Applied Photophysics, UK) in a 1 mm cuvette. The protein was heated from 20° C to 90° C at a rate of 1° C min^-1^ while ellipticity was monitored at 220 nm. The thermally induced unfolding of A20 OTU domains was not reversible. The unfolding of the A20 OTU domains was described by a two-state model^76^ (**Equation 1**).

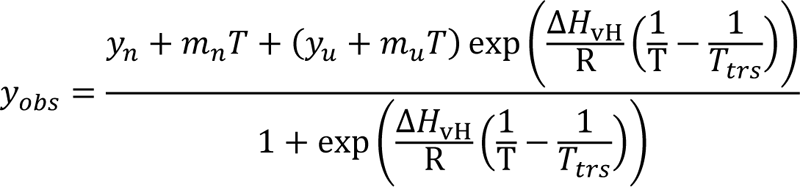

Where *y_obs_*is the observed elipticity, *y_n_*and *y_u_* are the elipticity values observed for the native and unfolded states, respectively. The *m_n_* and *m_u_* values are the linear temperature dependencies of *y_n_*and *y_u_*. Δ*H_vH_* is the apparent van’t Hoff enthalpy, R is the universal gas constant, and *T_trs_* is the temperature at which the population of unfolded protein is 50%. Curves were fit by non-linear regression using GraphPad Prism v6 (GraphPad Software).

### Statistical methods

Results are expressed as mean +/- standard error mean (SEM). Statistical analysis was performed using the Student’s *t*-test or ANOVA were indicated.

## Extended Data Figures and Tables

**Extended Data Fig. 1.**
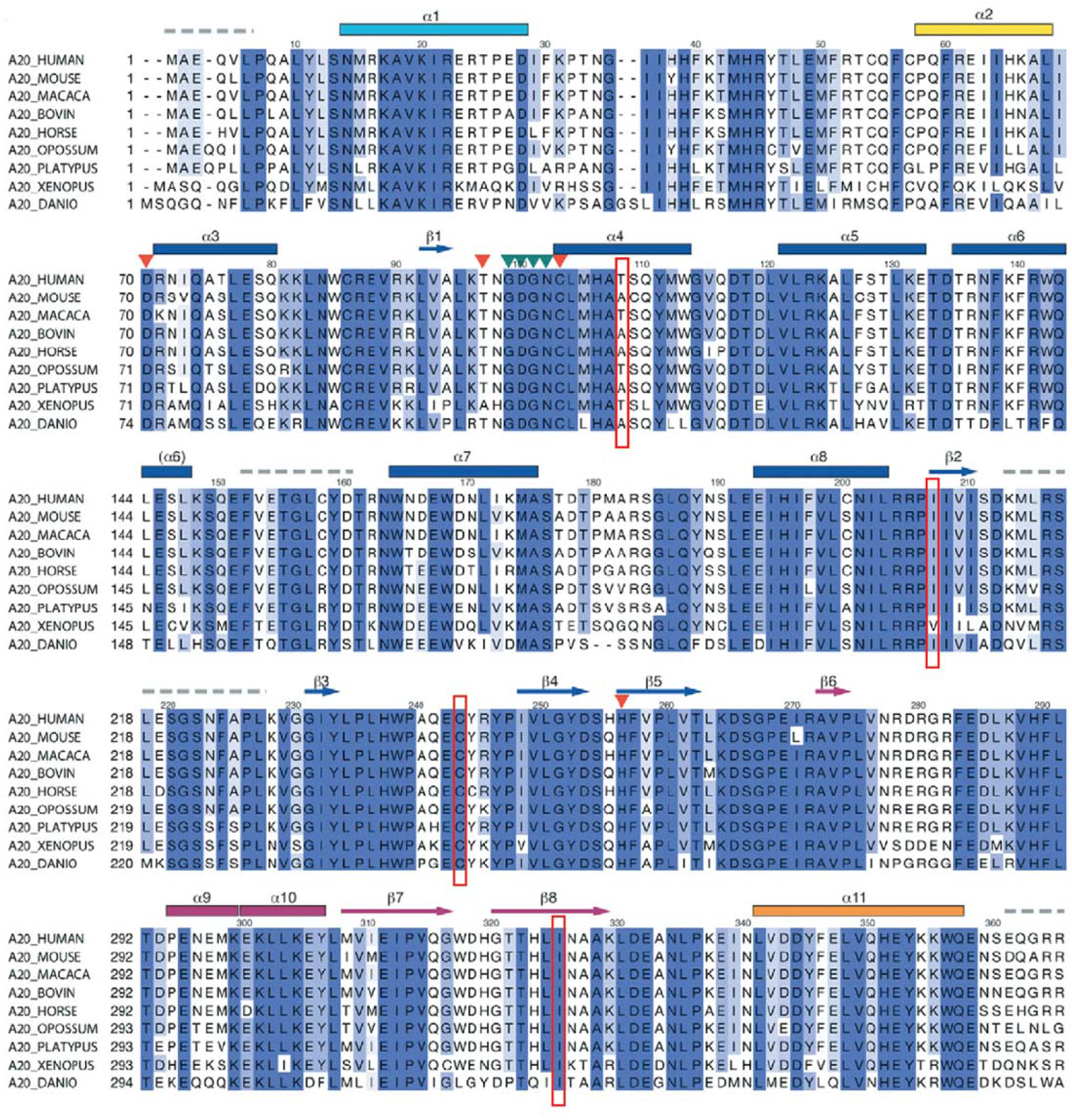
Evolutionary conservation of I207, C243 and I325 positions in A20 protein amongst jawed vertebrates. Aligned amino acid sequence and secondary structure elements of the A20 OTU domain, from ^40^. T108, I207, C243 and I325 residues are boxed in red.

**Extended Data Fig. 2.**
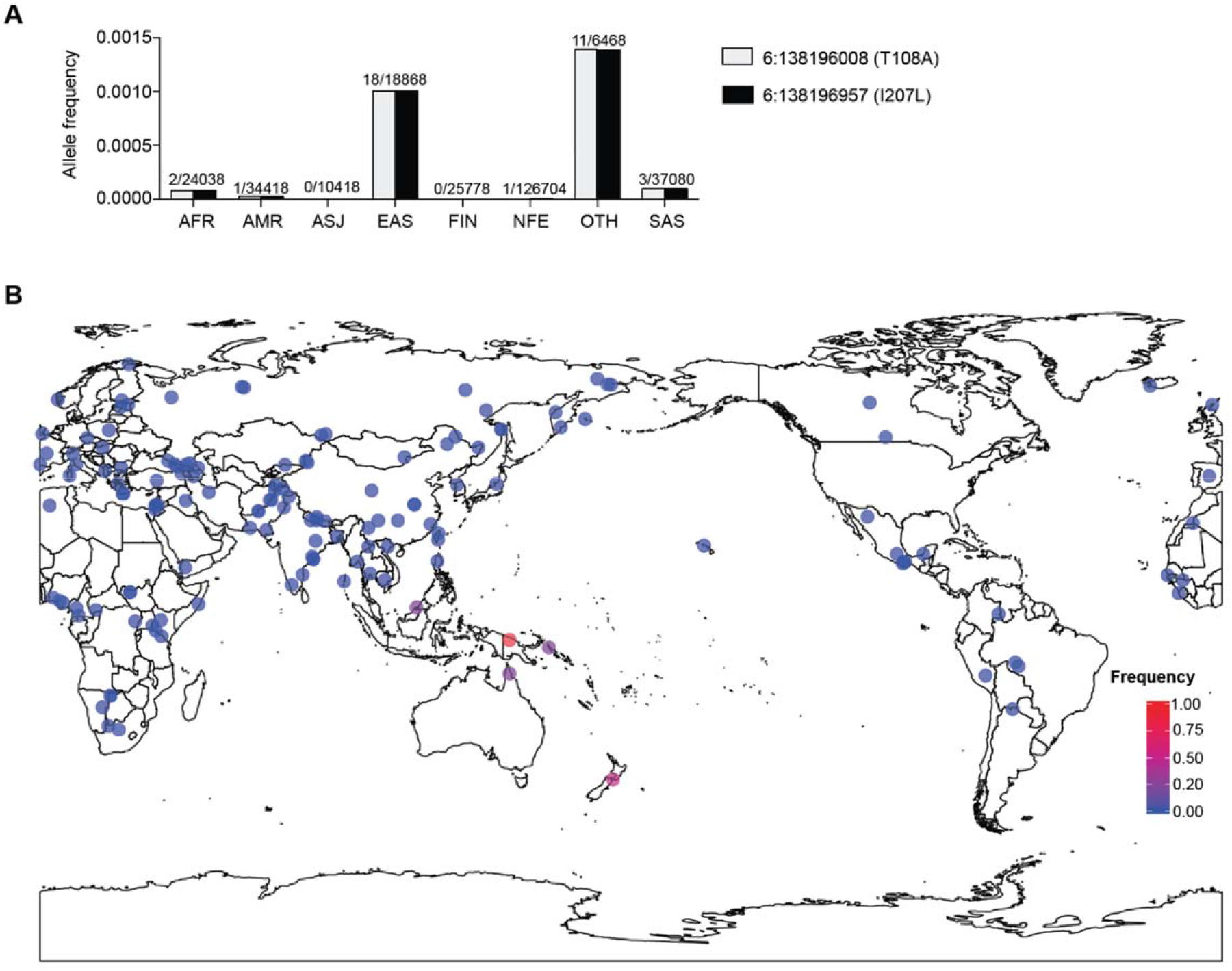
Global distribution of the T108A;I207L *TNFAIP3* haplotype. **(A)** gnomAD r2.0.2 population allele frequencies for the T108A and I207L missense variants in *TNFAIP3*, with allele counts above each bar (total as denominator). Populations are defined by principal component clustering of samples with individuals of known ancestry: those which do not cluster fall into the OTH group. AFR, African; AMR, Latino; ASJ, Ashkenazi Jew; EAS, East Asian; FIN, Finnish European; NFE, non-Finnish European; OTH, other; SAS, South Asian. The OTH population includes one homozygote, while all other alleles are heterozygous. **(B)** Frequency of the T108A;I207L *TNFAIP3* haplotype within 279 individual genomes from the Simons Genome Diversity Project^22^.

**Extended Data Fig. 3.**
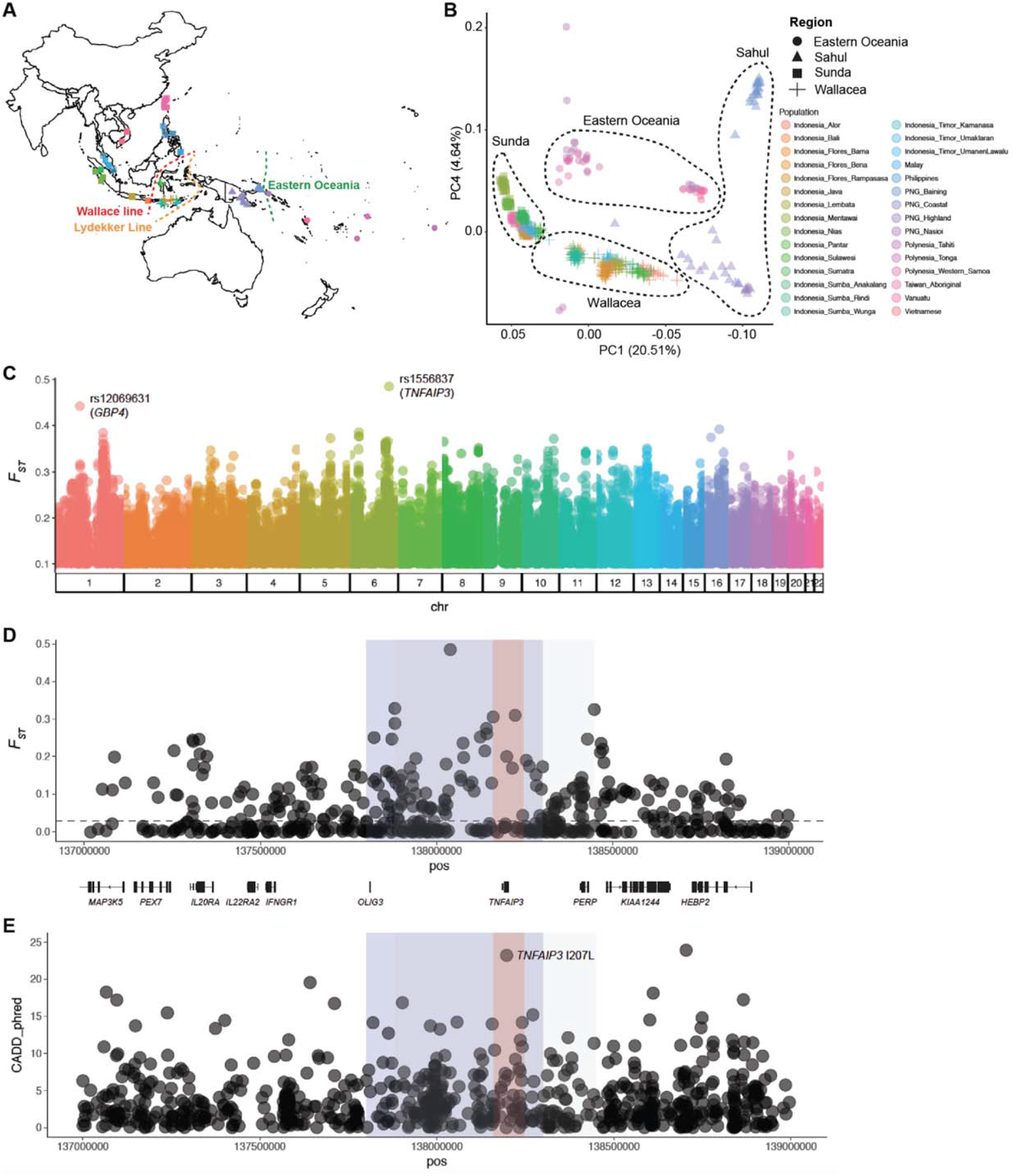
Population genetics of the Denisovan *TNFAIP3* haplotype. **(A)** Geographic distribution of samples used for population genetic analyses. Wallace Line (red) ^31^, Lydekker Line (yellow). Eastern Oceania is defined here as islands east of the Bismarck Archipelago (green line). **(B)** Principal component analysis (PCA), with the proportion of variance explained by each PC indicated on each axis **(C)** Genome-wide F_ST_ values between indigenous populations east and west of the Wallace Line. **(D)** Maximum F_ST_ values on chromosome 6 scores span the *TNFAIP3* locus. Shaded regions correspond to peak F_ST_ values between populations east and west of the Wallace Line (grey, 6:137881500-138448062), a previously described haplotype of elevated Denisovan ancestry in Oceanians (blue, 6:137800000-138300000, ^25^), and a putatively adaptively introgressed haplotype in Papuans (red, 6:138160925-138246514, ^29^). **(E)** Phred-scaled CADD scores of all PASS variants with gnomAD allele frequency <0.01 across the extended Denisovan haplotype ^27^. Shading as per panel D.

**Extended Data Fig. 4.**
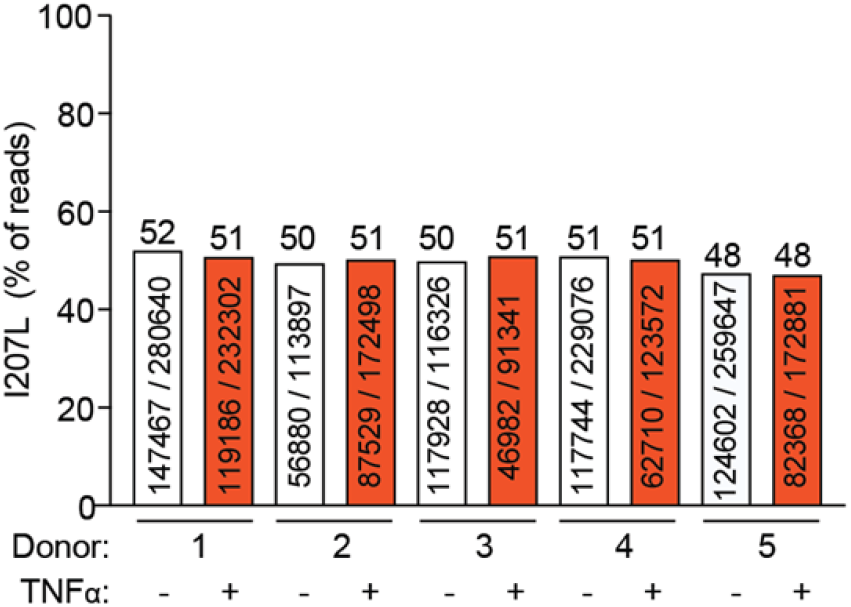
Relative abundance of Denisovan and modern human *TNFAIP3* mRNA from heterozygous leukocytes. Peripheral blood mononuclear cells from five healthy T108A;I207L heterozygous donors were cultured with (+) or without (-) TNFα for 2 hours, mRNA isolated and converted to cDNA and amplified with primers in exon 3 and 4 on either side of the I207 codon. The products were deep sequenced on an Illumina MiSeq. Shown are the fraction and percentage of reads derived from the T108A;I207L Denisovan allele.

**Extended Data Fig. 5.**
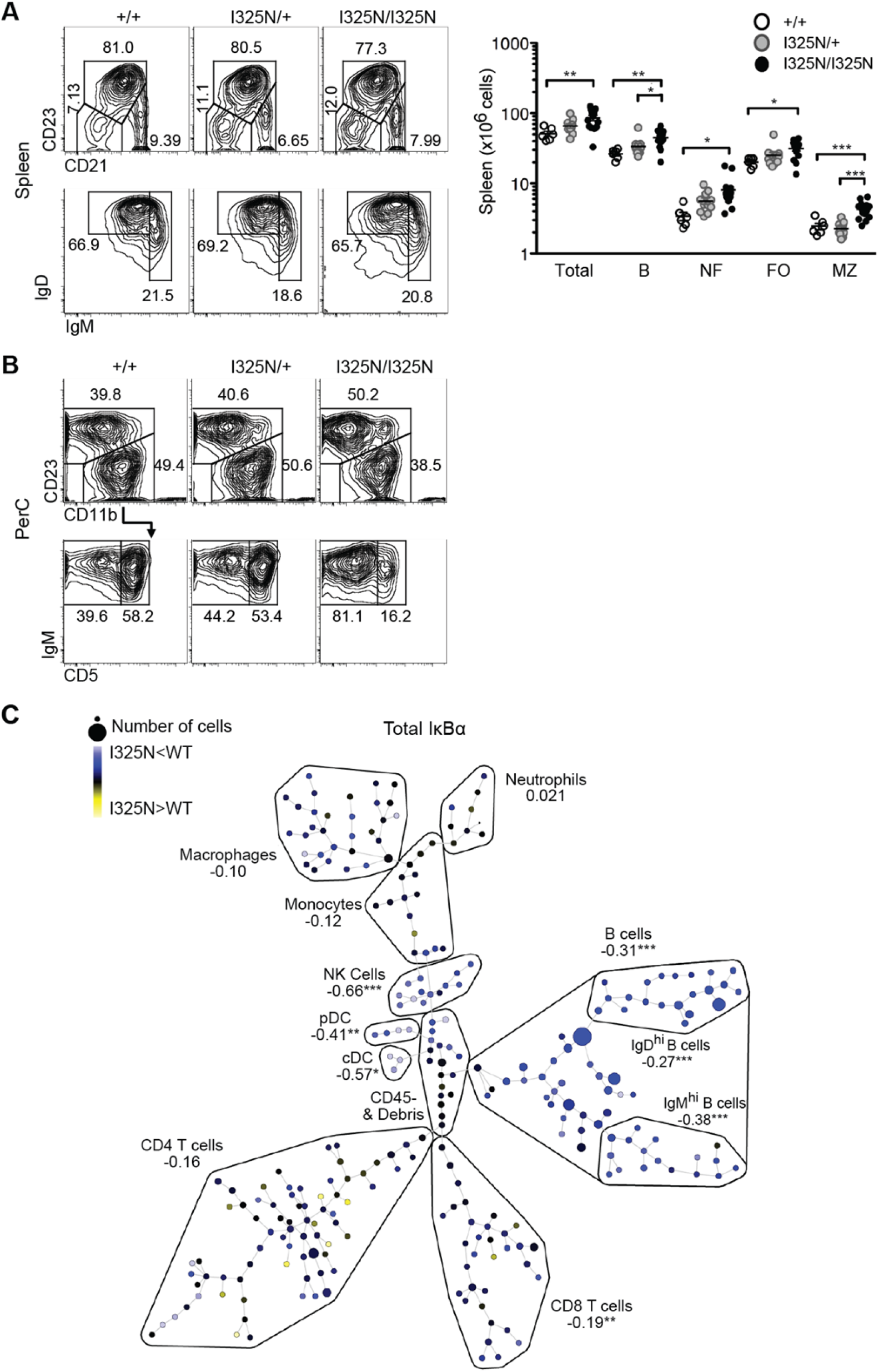
B cell lymphocytosis induced by the *Tnfaip3^I325N^* mutation. **(A)** Representative flow cytometry plots from mice of the indicated genotypes showing total frequency of splenic CD23^+^ and CD21^+^ positive cells (top panel). The bottom panel shows cells subsetted into IgD^+^ and IgM^+^ B cells. Note that the number of splenocyte B cell subsets is increased in *Tnfaip3*^I325N/I325N^ mice, shown to the right. **(B)** Representative flow cytometry plots from mice of the indicated genotypes showing the frequency of CD23^+^ and CD21^+^ positive cells in the peritoneal cavity (top panel). The bottom panel shows frequency of IgM^+^ and CD45^+^ B cells. Number of peritoneal cavity B cell subsets in mice of indicated genotypes is shown in Fig. 2B. **(C)** CYTOF analysis of intracellular IκBα in immune cells. Unstimulated spleen cells from *Tnfaip3*^+/+^ or *Tnfaip3*^I325N/I325N^ mice (*n*=4 per genotype) were individually labeled with mass-barcodes, mixed, permeabilized and stained with mass-labeled antibodies to a panel of cell surface markers and intracellular proteins including IκBα, and analyzed by CYTOF mass spectrometry^59,60^. Spanning-tree Progression Analysis of Density-normalized Events (SPADE; ^61^) analysis was used to resolve leukocyte lineages and subsets, and the relative intensity of IκBα in each subset was compared between *Tnfaip3*^I325N/I325N^ and wild-type cell counterparts. Shown by color and numbers is the mean hyperbolic arcsine (arcsinh) ratio in minor and major leukocyte subsets. Note that IκBα levels are reduced in B cell subsets indicating increased NF-κB activation. Significant differences indicated by Student’s *t*-test comparison of major subsets are marked: \**P* < 0.05; \*\**P* < 0.01; \*\*\**P* < 0.001.

**Extended Data Fig. 6.**
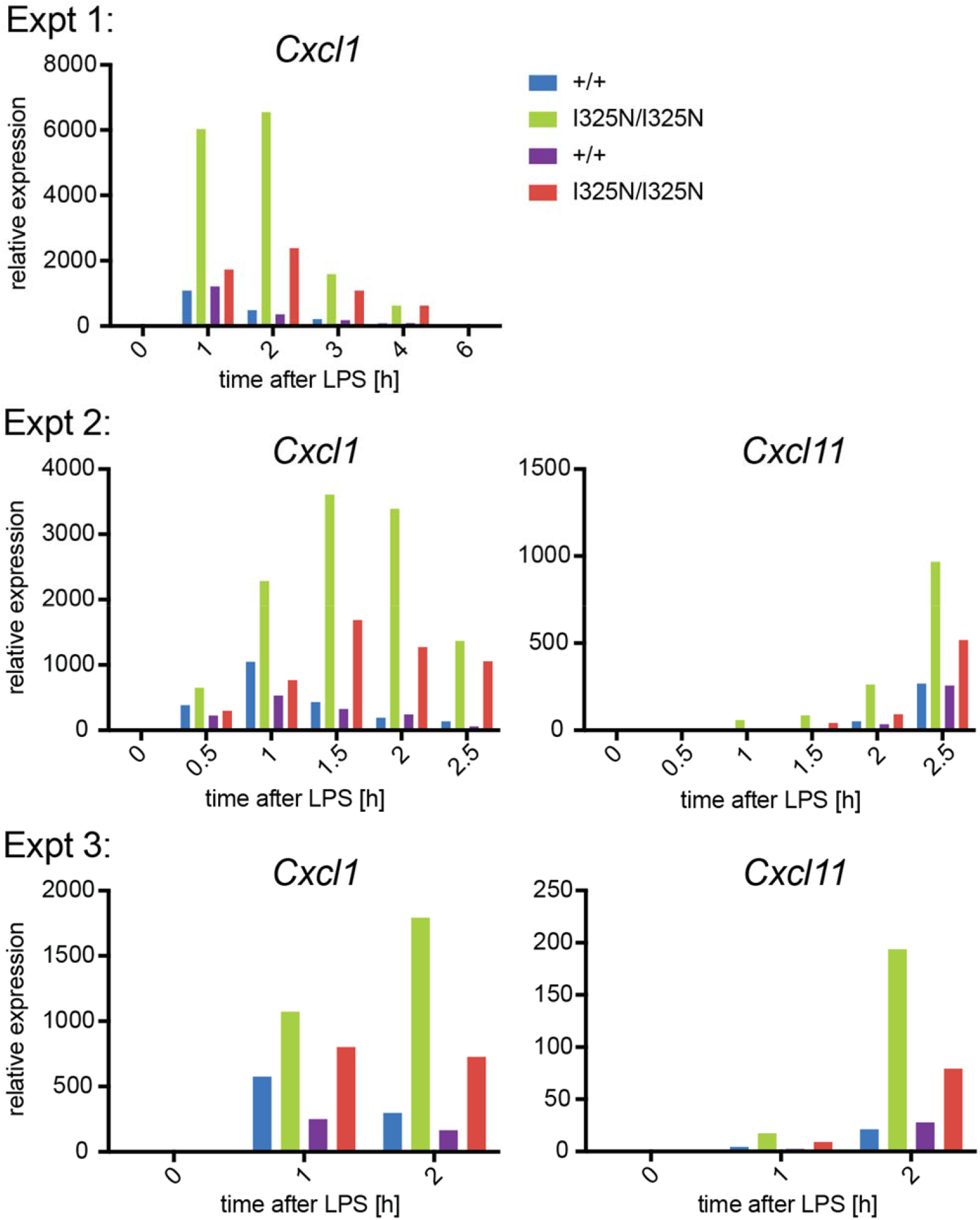
Exaggerated cytokine production by bone marrow-derived macrophages with the *Tnfaip3*^I325N^ mutation. Bone marrow macrophages derived from conditionally *Hoxb8*- immortalized bone marrow progenitor cells were derived from two separate pairs of wild-type and I325N homozygous mice, stimulated with LPS (10ng/ml) for the indicated times, and *Cxcl1* or *Cxcl11* mRNA determined by qPCR relative to *Ef1a* expression. Data are shown from three independent experiments, each with two different mouse donors of each genotype.

**Extended Data Fig. 7.**
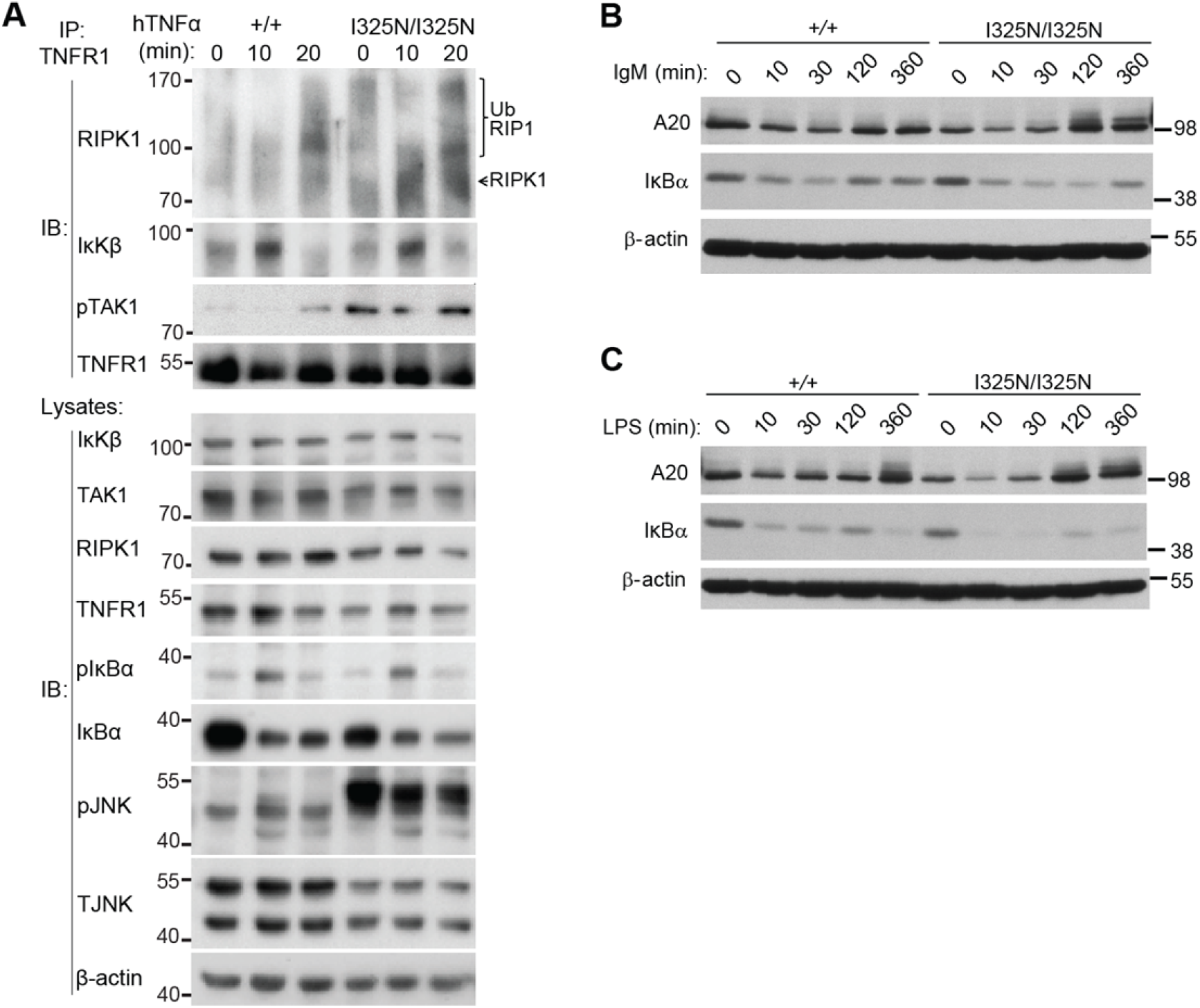
Prolonged NF-κB signaling in thymocytes and B cells extracted from I325N variant C57BL/6 mice. **(A)** Representative immunoblot analysis (IB; *n*=3 independent experiments) of canonical NF-κB components following immunoprecipitation (IP) of TNFR1 from lysates of *Tnfaip3*^+/+^ or *Tnfaip3*^I325N/I325N^ thymocytes treated with 200 U/ml hTNFα for the indicated times. Following TNFα stimulation RIPK1 was recruited to the TNFR1 complex, with increased high molecular weight polyubiquitinated forms of RIPK1 (UbRIP1) and auto-phosphorylated transforming growth factor β activated kinase-1 [TAK1^78^] in A20^I325N/I325N^ thymocytes (A). Mutant thymocytes exhibited a modest increase in IκBα degradation relative to wild-type, and increased JNK phosphorylation (Lysate blots). A similar pattern of enhanced TAK1 activation and JNK phosphorylation was seen in murine embryonic fibroblasts and bone marrow derived macrophages expressing catalytically dead A20 OTU or ZnF4 mutants^18^. **(B, C)** Wild-type (+/+) and I325N homozygous (I325N/I325N) *Tnfaip3* splenic B lymphocytes were stimulated for the indicated times with anti-IgM (B), or lipopolysaccharide (LPS) (C) and lysates analyzed by immunoblotting for A20, IκBα and β-actin (loading control). As observed for mutant thymocytes, B cells exhibited a modest increase in IκBα degradation relative to wild-type.

**Extended Data Fig. 8.**
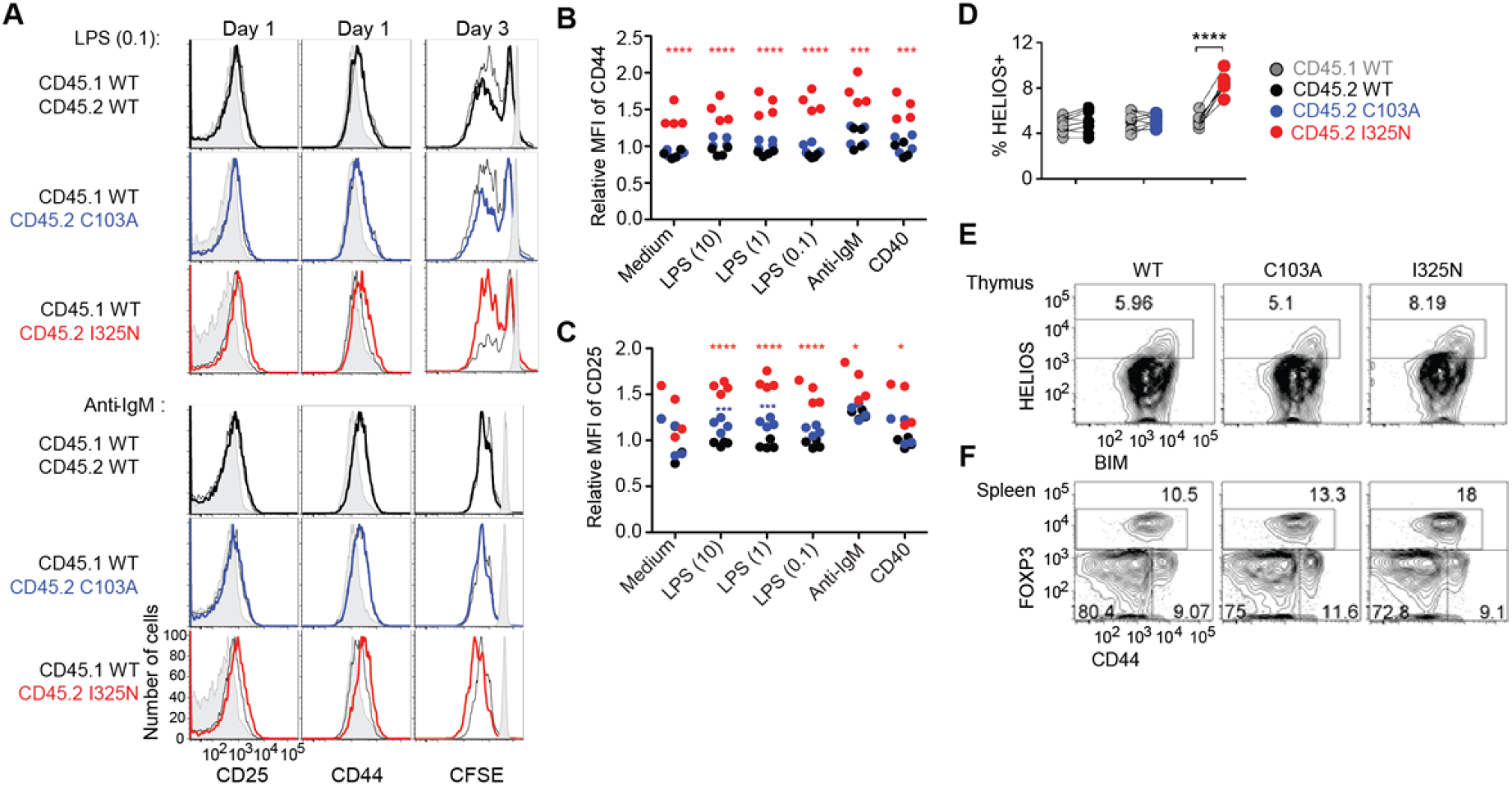
Cell autonomous exaggeration of B cell activation and Treg formation by Tnfaip3 I325N more than C103A mutations. B6.CD45.1^+^ wild-type mice were transplanted with a congenic bone marrow mixture from *Tnfaip3^+/+^* CD45.1^+^ donors, to provide wild-type lymphocytes as an internal control, and CD45.2^+^ donors of *Tnfaip3^I325N/I325N^*, *Tnfaip3^C103A/C103A^* or *Tnfaip3^+/+^* genotypes. **(A)** Representative flow cytometric histograms of CD25 and CD44 expression on B cells from chimeric mice cultured for 1 day and CFSE dilution after 3 days with 0.1 ug/ml LPS or anti-IgM. Black or colored histograms, CD45.2^+^ B cells of the indicated *Tnfaip3* genotype; grey line unfilled histograms, CD45.1^+^ control B cells in the same stimulated culture; grey filled histograms, B cells in a parallel unstimulated culture. **(B, C)** Data from independent mixed chimeric animals showing relative mean fluorescence intensity (MFI) of CD25 or CD44 on CD45.2^+^ B cells of the indicated genotypes (red, I325N; blue, C103A; black, WT) compared to the co-cultured CD45.1^+^ wild-type B cells. Statistical analysis by ANOVA: \**P* < 0.05; \*\**P* < 0.01; \*\*\**P* < 0.001; \*\*\*\**P* < 0.0001. Figure 2C shows the percentage of CD45.2^+^ cells of the indicated genotypes among viable B cells, relative to starting percentage, in cultures from individual mixed chimera donors stimulated with 0.1 ug/ml LPS. **(D)** Pairwise comparison of percent HELIOS^+^ cells among the indicated CD45.2^+^ and CD45.1^+^ subsets of CD4^+^ CD8^-^ CCR7^+^ CD24^+^ FOXP3^-^ thymocytes from the same chimera. Analysis by paired Student’s *t*-test; \*\*\*\**P* < 0.0001. Compared to the C103A mutation, the I325N mutation exaggerated thymic formation of FOXP3^+^ CD4^+^ cells and their Helios^+^ FOXP3^-^ precursors, which depend on TCR signaling through CARD11 to NF-κB. **(E, F)** Representative profiles gated on CD45.2^+^ cells of the indicated *Tnfaip3* genotypes. (E) Analysis of CD4^+^ CD8^-^ CCR7^+^ CD24^+^ FOXP3^-^ thymocytes, showing the percentage of immature medullary CD4 T cells induced by strong self-reactivity to express high levels of HELIOS and BIM. (F) Percentage of FOXP3^+^ CD44^+^ cells among CD4^+^ splenic T cells.

**Extended Data Fig. 9.**
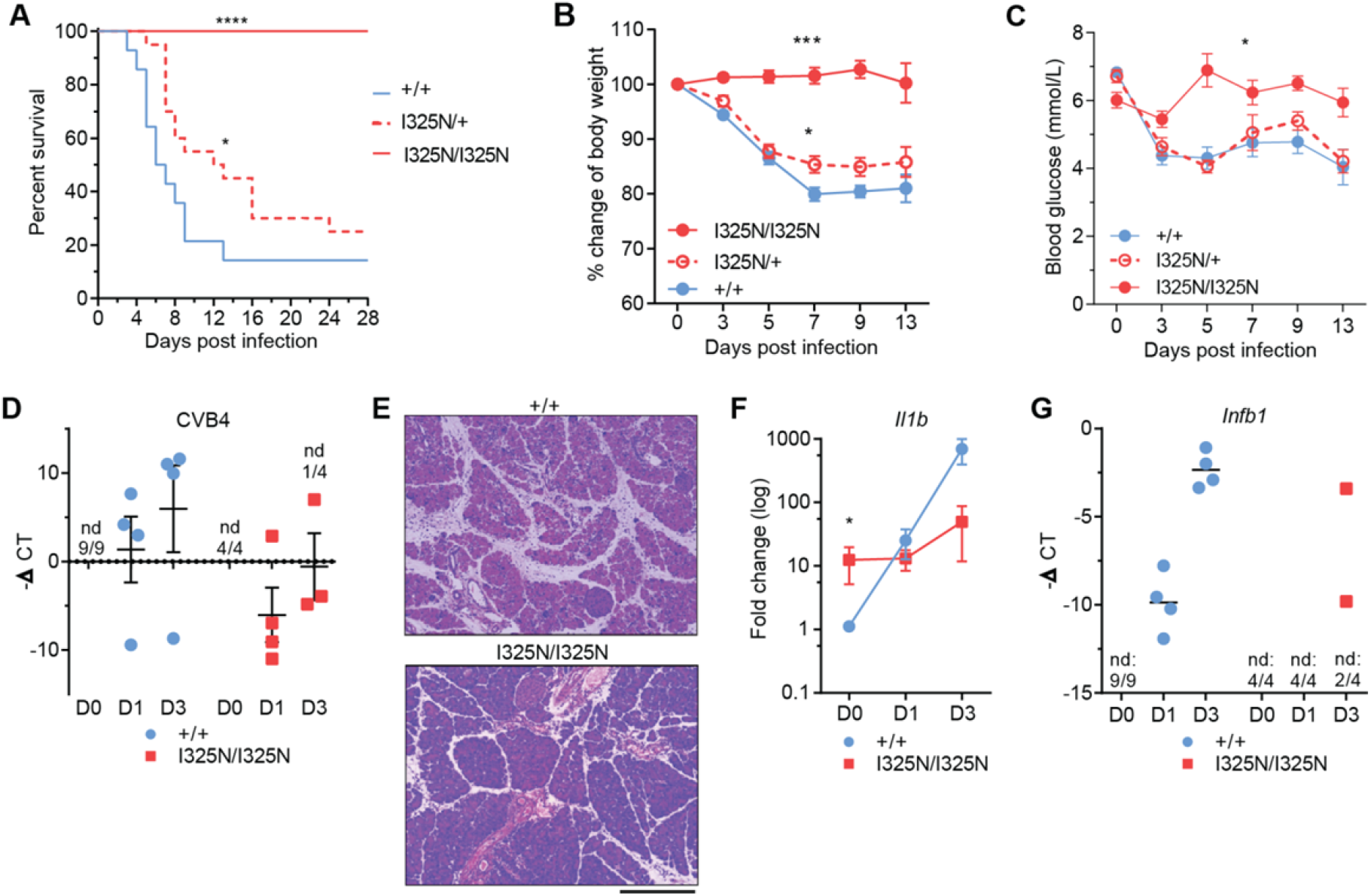
Increased resistance to Coxsackievirus B4 in A20 I325N mice. Infection of C57BL/6 male mice of the indicated *Tnfaip3* genotypes with 20 plaque forming units (PFU) of Coxsackievirus B4 (CVB4) E2 strain by intraperitoneal injection. **(A)** Kaplan-Meier survival curve showing a mean survival time of 6.5 and 12.5 days for *Tnfaip3*^+/+^ and *Tnfaip3*^+/I325N^ mice, respectively. No *Tnfaip3*^I325N/I325N^ mice succumbed to infection. Significance determined by Log-rank test. **(B)** Percent change in body weight after CVB4 infection and **(C)** Blood glucose levels of mice with indicated *Tnfaip3* genotype, compared to wild-type by area under the curve analysis. **(D)** CVB4 mRNA abundance on indicated days (D0-3) post infection. **(E)** Hematoxylin & eosin stained sections of pancreas from *Tnfaip3*^+/+^ and *Tnfaip3*^I325N/I325N^ mice at post-infection day 9. Note better preserved pancreatic architecture in *Tnfaip3*^I325N/I325N^ mice. Scale bar = 200 µm. **(F, G)** RTPCR analysis for inflammatory genes *Il1b* (F) or *Infb1* (G) at post-infection day 0, 1 and 3. Error bars represent SEM and Student *t*- test used for significance analysis unless otherwise stated, \**P* < 0.05; **\*\*\****P* < 0.001.

**Extended Data Fig. 10.**
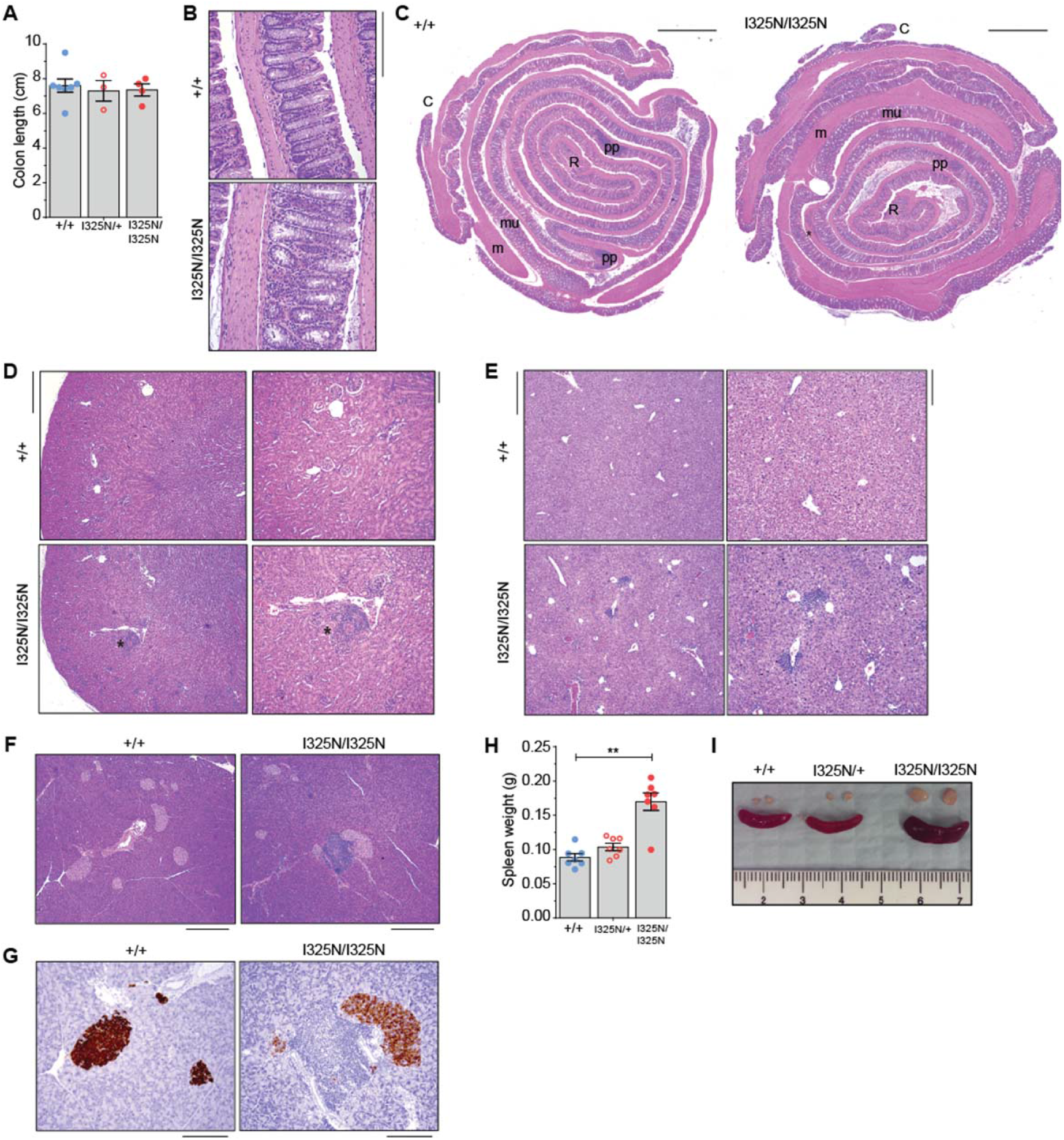
Histological analysis of tissues from A20^I325N^ mice. **(A)** Colon length (cm) and **(B)** Hematoxylin and eosin (H&E) stained sections from *Tnfaip3*^+/+^ and *Tnfaip3*^I325N/I325N^ mice colon (20 × magnification; scale bar = 200 µm) and **(C)** power-mosaic image of whole colon cross section (R=rectum; C=Caecum; m=muscularis; mu=mucosa; pp=Peyer’s patches; scale = 1 mm). **(D)** H&E sections of kidney (4 × left images, scale bar = 500 µm; 10 × right images, scale bar = 200 µm) and **(E)** liver (4 × left images, scale bar = 500 µm; 10 × right images, scale bar = 200 µm). **(F)** H&E stained sections of pancreas (10 X images, scale bar = 200 µm) or, **(G)** insulin stained sections (20 X images, scale bar = 100 µm). **(H)** Weight of spleens in grams (g). **(I)** Gross appearance of spleen and salivary glands from *Tnfaip3*^+/+^ and *Tnfaip3*^I325N/I325N^ mice. Images are representative from *n*=4 *Tnfaip3*^+/+^ and *n*=4 *Tnfaip3*^I325N/I325N^ 16 week old mice. Error bars represent SEM, \*\**P* < 0.01.

**Extended Data Fig. 11.**
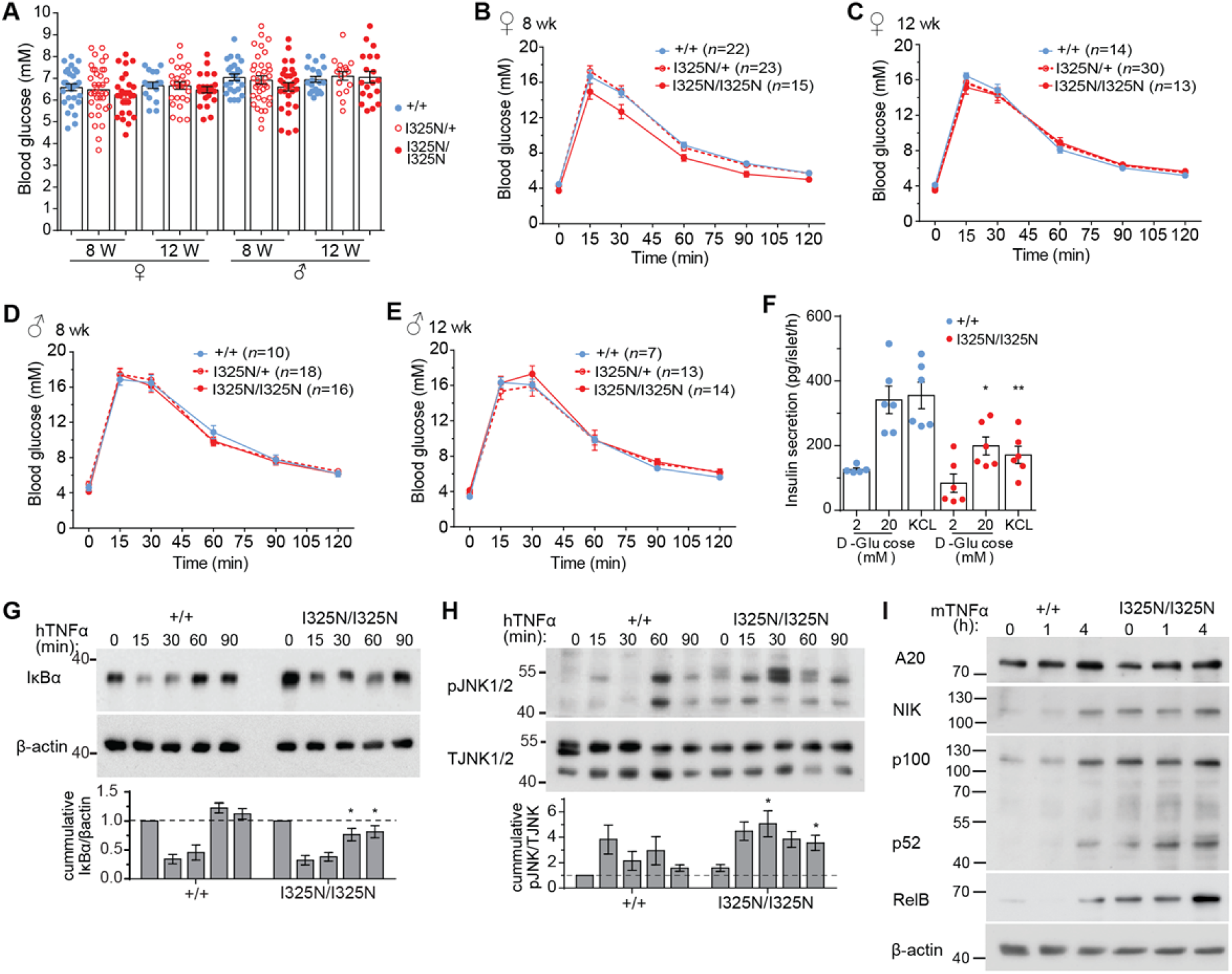
Glucose homeostasis of A20^I325N^ mice and isolated islets. **(A)** Random blood glucose levels of *Tnfaip3*^+/+^, *Tnfaip3*^I325N/+^ or *Tnfaip3*^I325N/I325N^ female (♀) or male (♂) mice at 8 or 12 weeks of age. **(B-E)** Blood glucose levels (mM) were monitored following an intraperitoneal injection of glucose (2 g/kg) in 8 (B, D) or 12 week-old (C, E) mice. **(F)** Pancreatic islets were isolated from individual mice of the indicated A20 genotypes and incubated overnight. Following incubation an *in vitro* Glucose-Stimulated Insulin Secretion (GSIS) response in conditions of 2 mM, 20 mM D-glucose, or 25 mM KCl was conducted in separate groups of islets. Error bars represent SEM and Student’s *t*-test used for significance between treatments, \**P* < 0.05; \*\**P* < 0.01. **(G-H)** Pancreatic islets were isolated from individual mice of the indicated A20 genotypes, incubated overnight, and treated with 200 U/ml TNFα for the indicated times. (G) Representative immunoblot for IκBα and β-actin (loading control) or (H) phosphorylated JNK (pJNK) and total JNK (TJNK, loading control). Cumulative densitometry from 5 independent experiments is shown below (*n*=5 *Tnfaip3*^+/+^ and *n*=6 *Tnfaip3*^I325N/I325N^ biological replicates). **(I)** Immunoblot for non-canonical NF-κB components NIK, p100/p52 and RelB; representative of 2 independent experiments. *Tnfaip3*^I325N/I325N^ islets exhibited increased activation of the non-canonical NF-κB pathway that can alter beta cell transcriptional programs to favor reduced insulin output ^38^.

**Extended Data Fig. 12.**
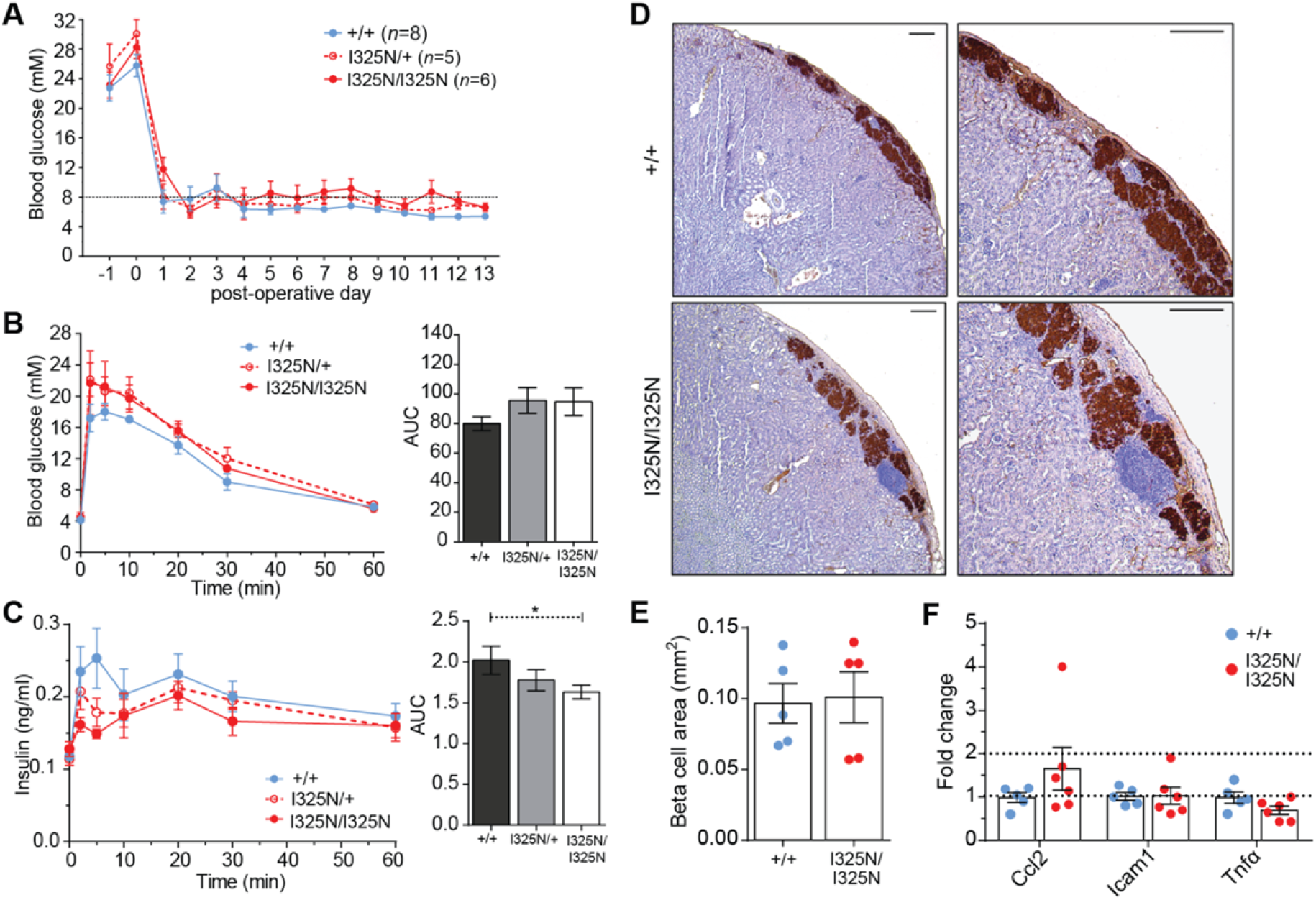
A20^I325N^ islet grafts exhibit a reduced first-phase insulin secretory response compared to wild-type A20 islet grafts independent to beta cell area. **(A)** Islets isolated from B6 mice of the indicated genotypes were transplanted under the kidney capsule of *Tnfaip3*^+/+^ B6 recipients that had been rendered diabetic with streptozotocin. Mean and SEM blood glucose on the indicated days relative to islet transplantation is shown. **(B-C)** At post-operative day 14 when euglycemia was established, mice were challenged with an (B) intravenous injection of glucose (1 g/kg), and blood glucose monitored over time (min). (C) Blood insulin levels (ng/ml) were also measured at the same time points via an enzyme-linked immunosorbent assay from samples in (B). AUC = area under the curve. Error bars represent SEM and ANOVA used for significance, \**P* < 0.05. **(D)** Insulin immunostaining of wild-type (*Tnfaip3*^+/+^) or A20 I325N homozygous mutant (*Tnfaip3*^I325N/I325N^) islet grafts transplanted under the kidney capsule of diabetic C57BL/6 mice for 30 days (*Tnfaip3*^+/+^ > *Tnfaip3*^+/+^ *n* = 5; *Tnfaip3*^I325N/I325N^ > *Tnfaip3*^+/+^ *n*=5). **(E)** Islet graft beta cell area was determined by insulin-positive area quantification in continuous serial graft sections. *Tnfaip3*^I325/I325N^ grafts exhibited equivalent insulin-positive graft area, confirming that loss of glucose tolerance (B, C) was due to a defect in insulin secretion. **(F)** *Tnfaip3*^+/+^ or *Tnfaip3*^I325N/I325N^ islet grafts from an independent cohort of recipients were isolated on post-operative day 10 and analyzed for the indicated mRNAs by RT-qPCR ^77^. The difference in gene expression of indicated islet grafts is shown. Fold change calculated using average wild-type ΔCt value.

**Extended Data Fig. 13.**
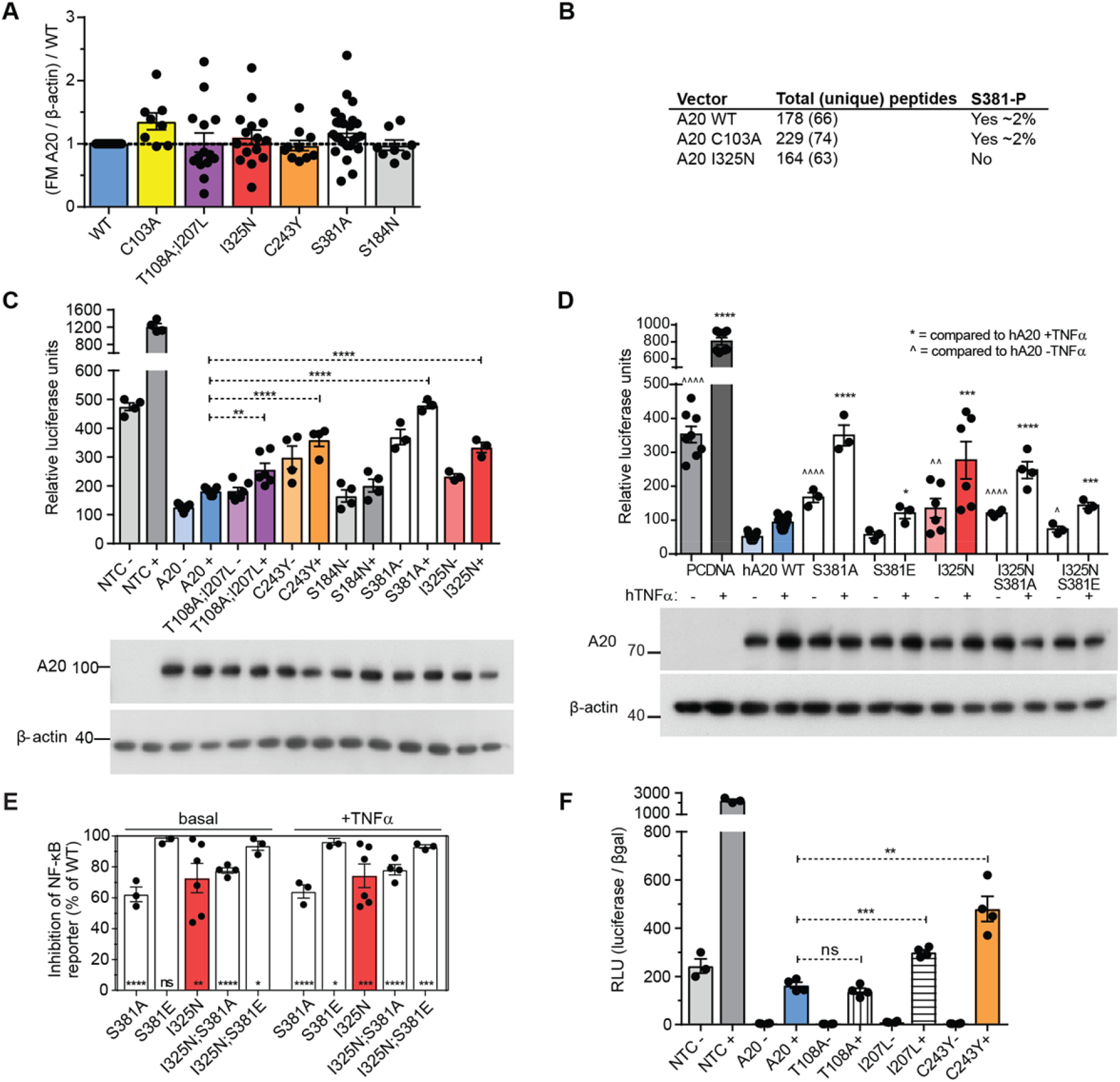
*In vitro* analysis of missense variant effects on A20 control of NF-κB. **(A)** A20 accumulation in βTC3 mouse insulinoma cells transfected with wild-type (WT) A20, C103A A20, T108A;I207L A20, I325N A20, C243Y A20, S381A A20 or S184N A20 was determined by densitometry analysis of experimental immunoblots. Fast migratory (A20) 98 kDa A20 was corrected to loading control and compared to WT A20 of the same immunoblot [(FM A20/β-actin)/A20]. **(B)** Results of mass spectrometry analysis of full length WT A20, C103A mutant A20, or I325N mutant A20 expressed in and purified from HEK293 cells and analysed by mass spectrometry for phosphorylation of the indicated residues as per references ^18,19^. **(C)** βTC3 cells co-transfected with an NF-κB.luciferase reporter and a CMV.βgal expression construct alone or with PCDNA3.1 encoding human wild-type A20 (hA20 WT; blue), A20 T108A/I207L (purple), A20 C243Y (orange), A20 S184N (grey), A20 I325N (red) or A20 S381A (white). Cells were stimulated with (+) 200 U/ml hTNFα for 8 h or left untreated (-). Data presented as relative luciferase units. Each column represents 3-5 independently transfected aliquots of cells within one experiment marked by circle symbols (representative of 3 independent experiments) and columns are arithmetic means. Statistical comparison by Student’s *t*-test against WT hA20 (blue). **(D-F)** βTC_3_ cells co-transfected with a NF-κB.luciferase reporter and a CMV.βgal expression construct alone or with PCDNA3.1 encoding human wild-type A20 (hA20 WT; blue), or (D) A20 I325N variant (I325N; red), or A20 with serine 381 substituted to non-phosphorylatable alanine (S381A or I325N S381A), or to phosphomimetic glutamate (S381E or I325N;S381E), (F) or A20 with the point variants T108A (vertical stripes), I207L (horizontal stripes) or C243Y (orange). Cells were stimulated with (+) 200 U/ml hTNFα for 8 h or left untreated (-). In (E) the two missense variants in the Denisovan haplotype, T108A and I207L, are tested individually (cross-hatched columns). Data presented as relative luciferase units. Each column represents mean of 3-6 independently transfected aliquots of cells within one experiment marked by circle symbols (representative of 3 independent experiments) and columns are arithmetic means. Statistical comparison by Student’s *t*-test against WT hA20 (blue) with TNFα (+) indicated by * or without (-) indicated by ^. In (F), data from (D) are presented as inhibition of NF-κB reporter as a percent of WT A20. Error bars represent SEM, \**P* < 0.05; \*\**P* < 0.01; \*\*\**P* < 0.001; \*\*\*\**P* < 0.0001.

**Extended Data Fig. 14.**
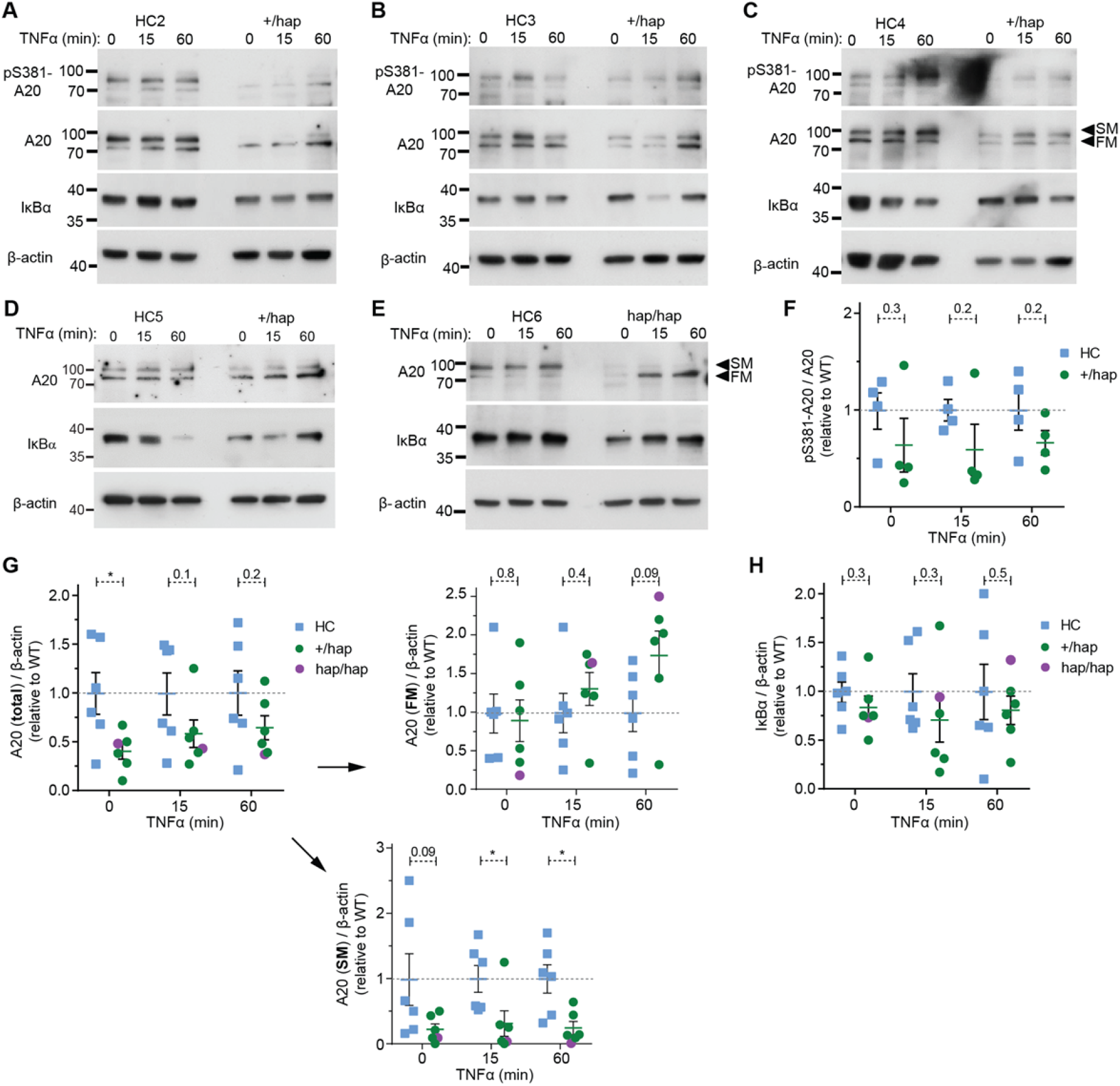
Immunoblot analysis of peripheral blood mononuclear cells from healthy individuals with the T108A I207L haplotype. **(A-E)** Immunoblot analysis of peripheral blood mononuclear cells (PBMC) from individuals with the Denisovan *TNFAIP3* haplotype (hap) or without (healthy control donor; HC). PBMCs were left untreated or stimulated with recombinant human TNFα (hTNFα) for 15 or 60 min. Proteins assessed include A20, phosphoserine-A20 (pS381-A20), IκBα and β-actin (loading control). **(F)** Densitometry analysis of pS381-A20 levels in immunoblots A-C and Fig. 4D. Densitometry was calculated by correcting to total A20 present and using the average value from samples without the T108A;I207L haplotype (HC) to compare all samples. **(G)** Densitometry analysis of total A20 levels in immunoblots A-C and Fig. 4D. Densitometry was calculated by correcting to β-actin loading control and using the average WT value to compare all samples. (G) is further divided into fast migrating (FM) or slow migrating (SM) A20. No significant difference is observed for FM A20 between hap or HC individuals. In contrast, a significant difference is observed between hap and HC individuals for SM A20, which correlates with phosphorylated A20 in *in vitro* studies (Fig. 4D, E) ^18,19^; consistent with reduced phosphorylation for A20 T108A;I2107L. **(H)** Densitometry analysis of IκBα levels in immunoblots A-C and Fig. 4D. Densitometry was calculated by correcting to β-actin loading control and using the average value from samples without the T108A;I207L haplotype (HC) to compare all samples. Each symbol represents an individual lane in immunoblot. Error bars represent SEM and Student’s *t*-test used for significance analysis unless otherwise stated, \**P* < 0.05.

**Extended Data Fig. 15.**
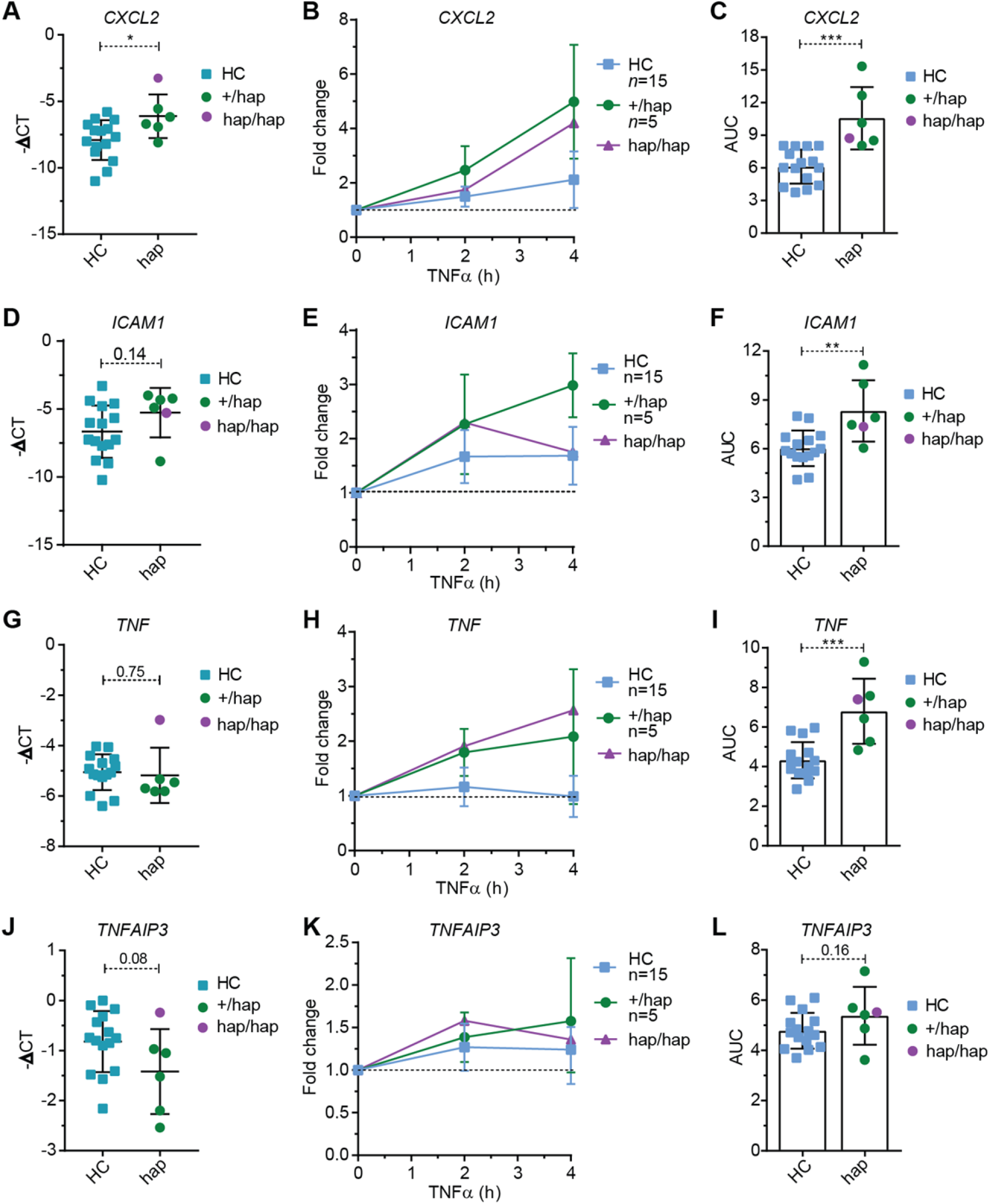
Peripheral blood mononuclear cells from individuals with the Denisovan *TNFAIP3* T108A;I207L haplotype are hyper-inflammatory. RT-PCR analysis of *CXCL2*, *ICAM1*, *TNF*, and *TNFAIP3* genes expression from peripheral blood mononuclear cells (PBMCs) from individuals heterozygous (green circles) or homozygous (purple circle) for the Denisovan *TNFAIP3* haplotype (hap) and 15 healthy control donors (HCs; blue squares), comparing negative ΔCT basal expression levels **(A, D, G, J)** or induction kinetics following stimulation with recombinant human TNFα for 0 2 or 4 h **(B, E, H, K).** Fold change was determined by comparing stimulated values against non-stimulated PBMC preparations within the same group. Significance between groups was determined by area under the curve analysis **(C, F, I, L)**. Error bars represent SD and Student’s *t*-test used for significance analysis unless otherwise stated, \**P* < 0.05; **\*\*\****P* < 0.001.

**Extended Data Fig. 16.**
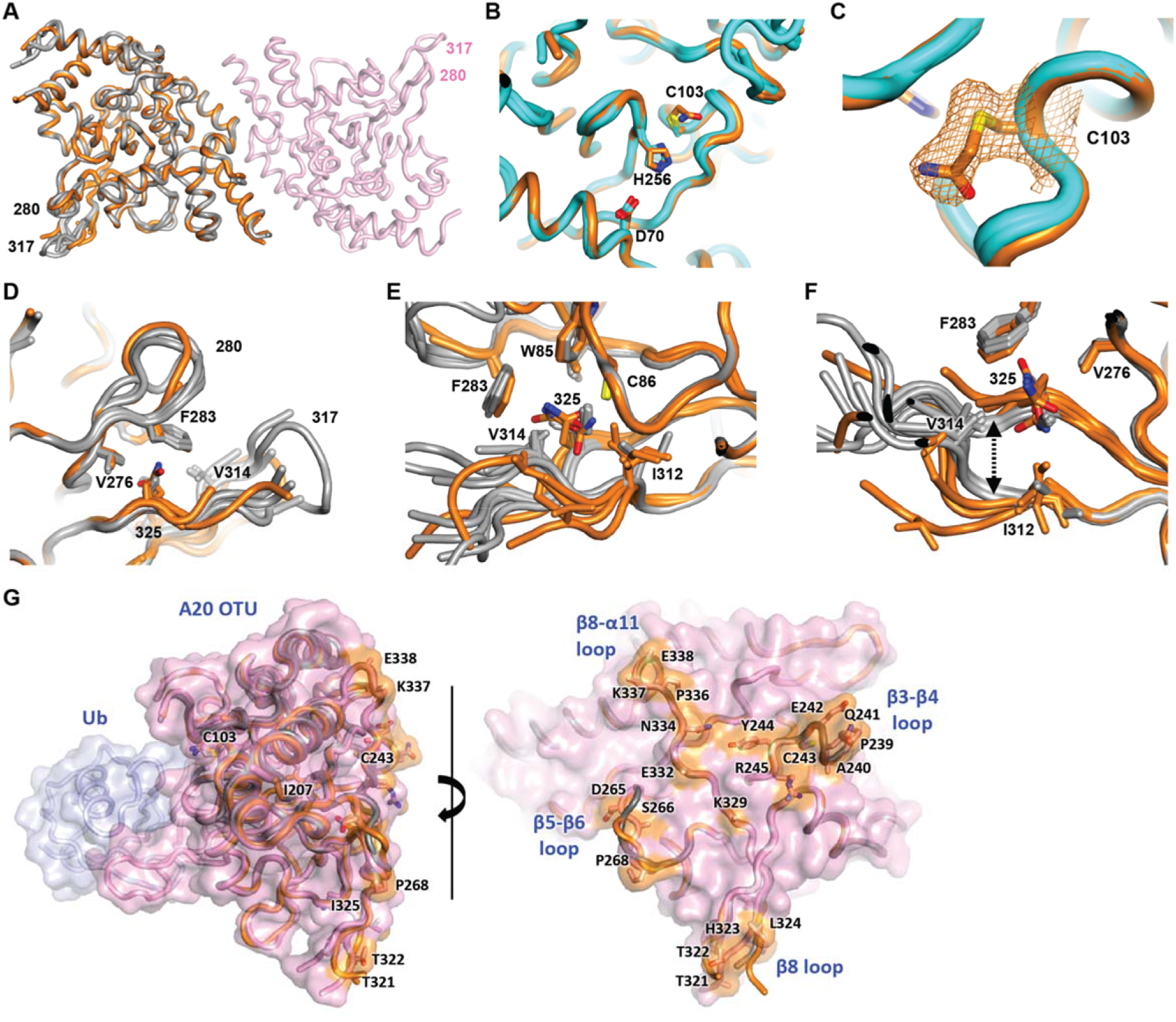
A20^I325N^ mutation subtly impacts the OTU structure. **(A)** Structures of human I325N variant (orange) and wild-type (WT; grey) A20 OTU domains superposed (each with 6 molecules per asymmetric unit). The dimer partner of one of these is shown in pink. Loop 317 and 280 are indicated on the periphery. **(B)** The I325N substitution does not alter the fold of the putative catalytic triad (orange cartoon and sticks) compared to previously published wild-type structures (cyan cartoon and sticks; PDB entries 3DKB, 2VFJ and 3ZJD). **(C)** The fold is additionally unperturbed by acetamidylation of C103 (orange mesh; 2mFo-DFc composite omit map contoured at 1.2 sigma). **(D-F)** The I325N substituted side-chain sits at the base of the β7-β8 loop containing residue 317, in a pocket also lined by the β6-α9 loop containing residue 280 (D). Adjacent hydrophobic residues include W85, V276, F283, I312, V314 (E). The I325N substitution results in the base of the β7-β8 loop splaying apart (F, arrow), increasing β7-β8 loop disorder. **(G)** Structures of human I325N variant (orange) and wild-type (WT; grey) A20 OTU domains superposed on the wild-type OTU-ubiquitin structure 5LRX ^42^. Conserved residues on the posterior surface are coloured orange. The β3-β4 loop containing the C243Y mutation and the structured base of the β7-β8 loop containing the I325N mutation are marked.

**Extended Data Fig. 17.**
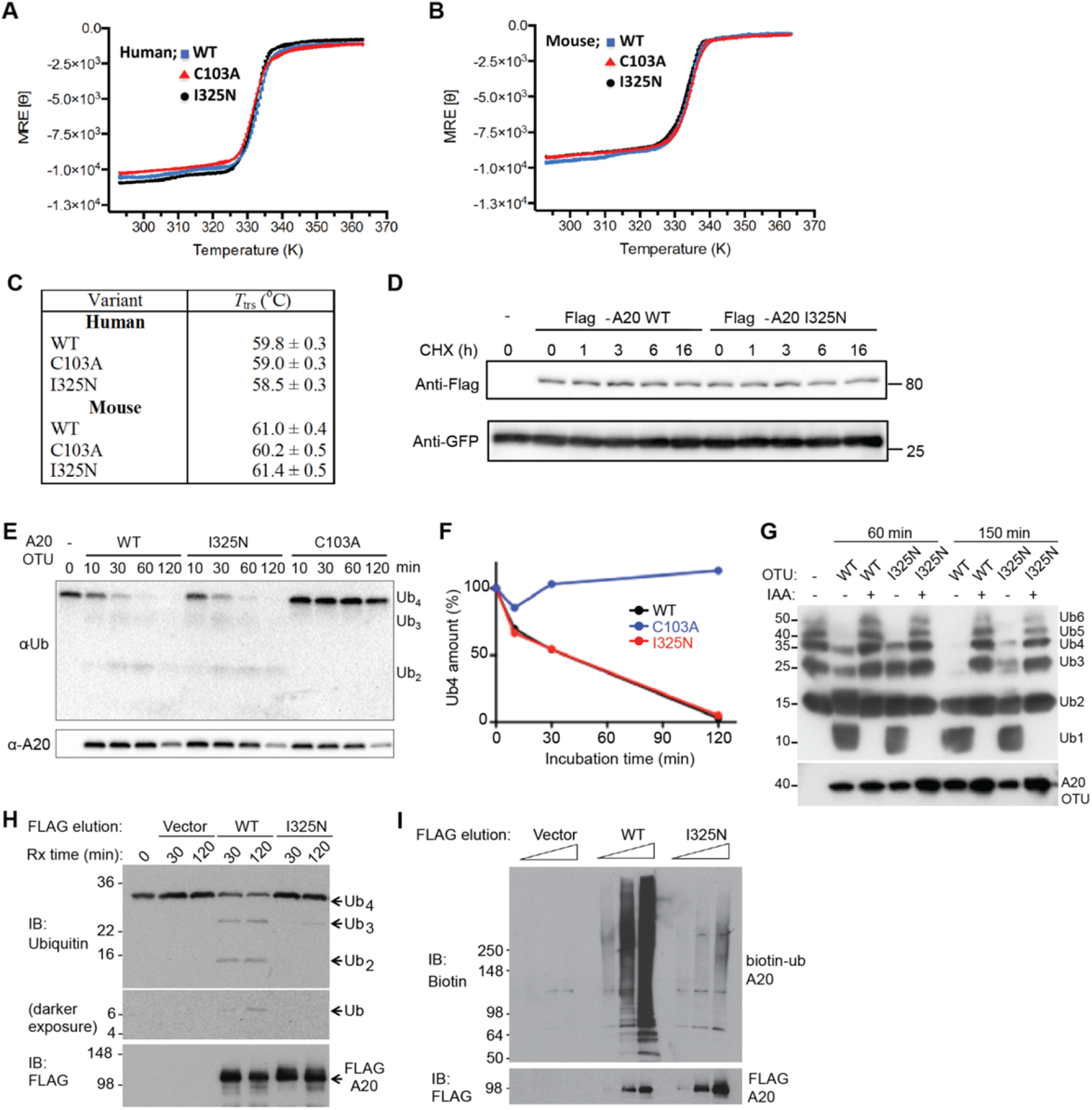
Molecular characterization of A20 I325N and A20 C103A. **(A-B)** Thermal stability of mouse and human WT, C103A, and I325N variants were examined using circular dichroism, where mean residue ellipticity (MRE) was measured at 220 nm as a function of temperature as solutions were heated at a rate of 1° C/min. Human (A) or mouse (B) A20 OTU residues 1- 370; WT (blue squares), C103A (red triangles), I325N (black circles). Curves were fit by linear regression. **(C)** Unfolding transitions temperature (T_trs_) of human A20 OTU variants (residues 1-370) and mouse variants (residues 1-360), derived from the fit of the curves in (A) and (B). **(D)** HEK293T cells were transfected with bicistronic retrovirus vector pMXs-IRES-GFP, expressing FLAG-tagged mouse full-length TNFAIP3 wild-type or I325N. Each group of transfected cells was replated 24 h after transfection and incubated with 10 µM of cycloheximide (CHX) for the indicated times before harvesting. Lysates were probed with anti-FLAG and anti-GFP antibodies. **(E, F)** Reactions containing K48-linked tetra-ubiquitin (Ub_4_) and *E. coli*-expressed and purified WT or I325N A20 OTU domain, or the catalytically deactivated A20 OTU domain, C103A, were incubated for the indicated times. Depolymerization of polyubiquitin was detected by immunoblotting for ubiquitin and A20; representative of two independent experiments. **(G)** Reactions containing mixed Ub_2_-Ub_6_ K48-linked polyubiquitin chains and *E. coli*-expressed and purified WT or I325N A20 OTU domain ± iodoacetamide (IA) treatment were incubated at 37°C for 60 or 150 minutes. Depolymerization of polyubiquitin was detected by immunoblotting for ubiquitin and the N-terminus of A20; representative of 4 independent experiments. **(H, I)** Flag-tagged full length human A20 proteins (WT or I325N mutant) were expressed in HEK-293T cells and purified as previously described ^14^. Purified A20 proteins were tested for K63-linked tetra-ubiquitin depolymerization (H) and *in vitro* ubiquitination assay (I).

**Extended Data Fig. 18.**
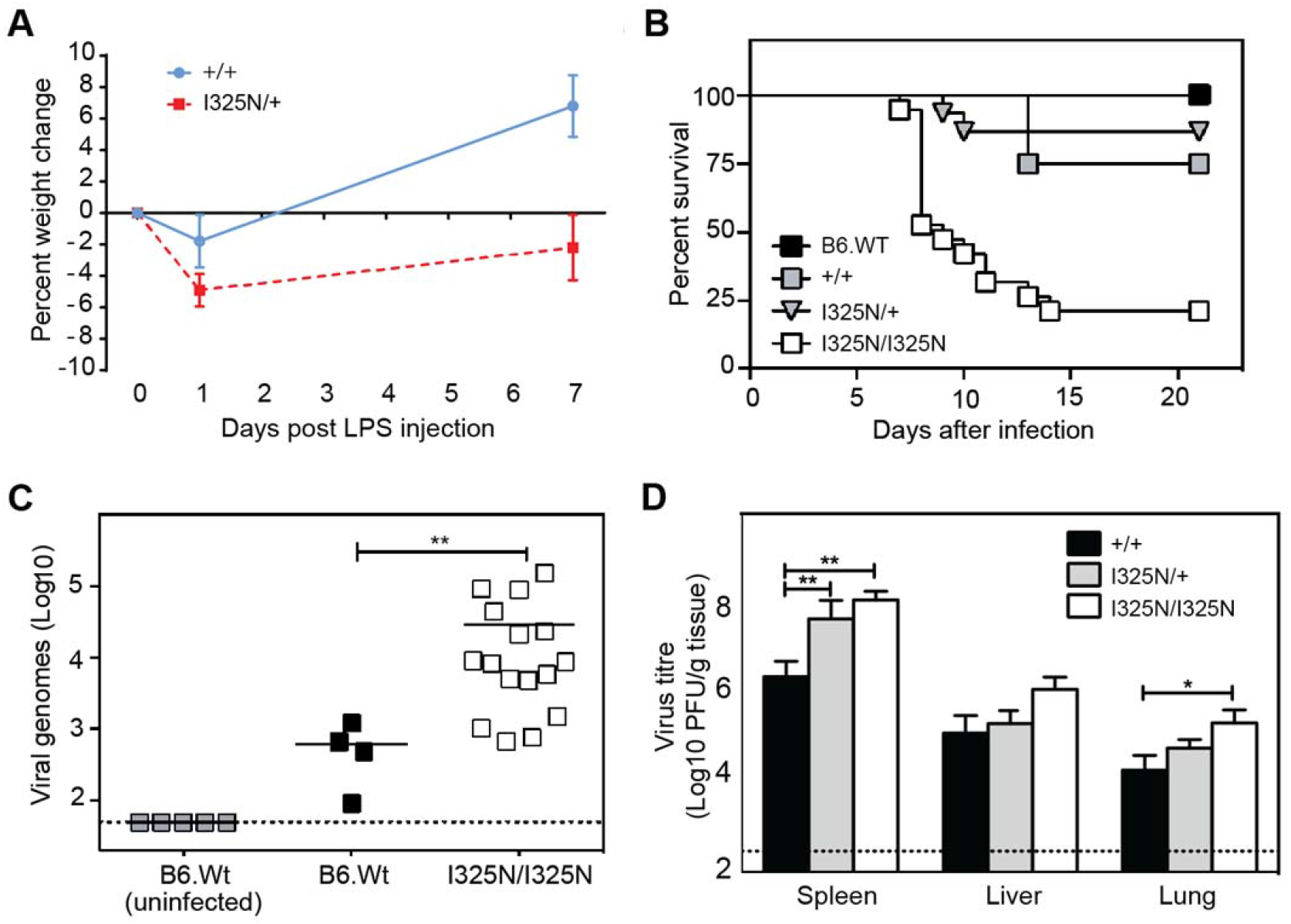
A20 I325N are susceptible to LPS injection and poxvirus infection. **(A)** Daily body weights for *Tnfaip3*^+/+^ or *Tnfaip3*^I325N/+^ mice administered 50 µg bacterial LPS by intraperitoneal injection. (B-D) Ectromelia virus infection of B6 male siblings of the indicated *Tnfaip3* genotypes and wild-type B6 control mice, showing **(B)** Kaplan-Meyer survival plots and **(C)** viral load in blood on day 8; \*\**P* < 0.005 by unpaired *t* test. **(D)** Virus load in spleen, liver and lung, respectively. Data expressed as mean ± SEM; data was log-transformed and statistical significance determined by 2-way ANOVA; \**P* < 0.05, \*\**P* < 0.01. Dotted lines in (C, D) denote limit of detection.

**Extended Data Fig. 19.**
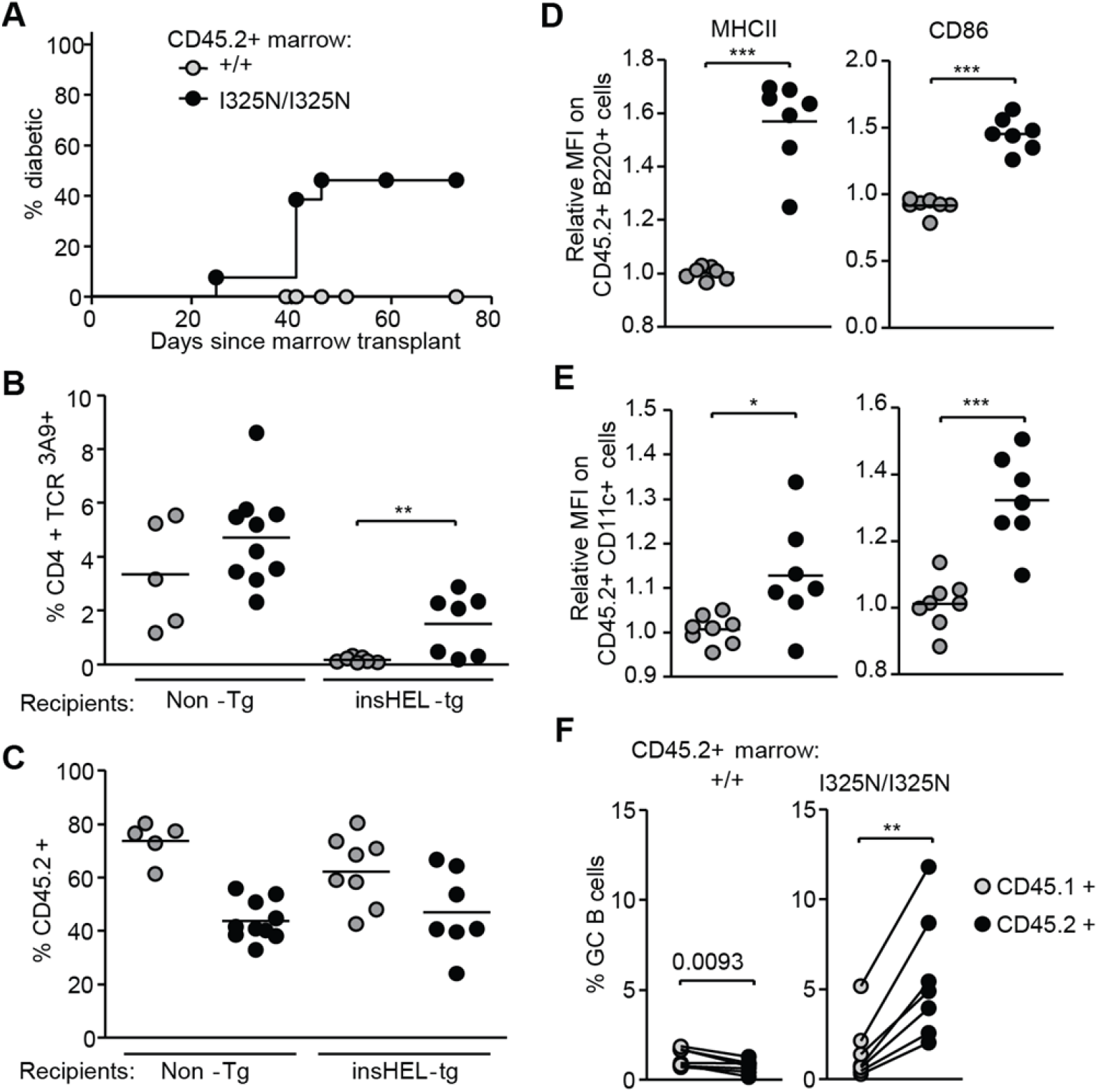
Diminished tolerance to self-antigen caused by mutant A20. InsHEL-transgenic mice or non-transgenic controls, both with wild-type A20 in pancreatic islets, were transplanted with a congenic bone marrow mixture from *Tnfaip3^+/+^* CD45.1^+^ TCR^3A9^-transgenic donors, to serve as an internal control, and CD45.2 TCR^3A9^-transgenic donors with either *Tnfaip3*^I325N/I325N^ (black circles) or *Tnfaip3^+/+^* (grey circles) genotype. **(A)** Incidence of diabetes in insHEL-transgenic recipients as a function of time after transplantation (*n*=12 chimeras for +/+ and 14 chimeras for I325N/I325N pooled from 2 separate experiments). **(B)** Frequency of TCR^3A9^ positive CD4^+^ T cells among CD45.2^+^ *Tnfaip3^+/+^* (grey circles) or *Tnfaip3*^I325N/I325N^ (black circles) lymphocytes in pancreatic lymph nodes. Test recipients expressed the insHEL antigen in pancreatic islets (insHEL-transgenic), while parallel non-transgenic recipients lacked the autoantigen. **(C)** Percentage of CD45.2^+^ *Tnfaip3^+/+^* (grey circles) or *Tnfaip3*^I325N/I325N^ (black circles) cells among TCR^3A9+^ CD4^+^ T cells in spleen. **(D-F)** Analysis of insHEL transgenic recipients. Relative mean fluorescence intensity (MFI) of staining for cell surface MHC class II or CD86 on CD45.2^+^ *Tnfaip3^+/+^* (grey circles) or *Tnfaip3*^I325N/I325N^ (black circles) B cells (D) or CD11c^+^ dendritic cells (E), compared to CD45.1^+^ *Tnfaip3^+/+^* B cells or dendritic cells in the same sample. **(F)** Frequency of germinal center (GC) B cells among CD45.2+ B cells (black circles) compared to the frequency among CD45.1^+^ *Tnfaip3^+/+^* B cells in the same animal (linked by lines). Left panel, CD45.2 *Tnfaip3^+/+^* marrow donor; right panel, CD45.2 *Tnfaip3*^I325N/I325N^ marrow donor. In (B-F), each symbol, or pair of symbols joined by a line, represents one chimeric mouse, all analyzed in a single experiment. Statistical analysis by 2-tailed Student’s *t*-tests: \**P* < 0.05; \*\**P* < 0.01; \*\*\**P* < 0.001.

**Extended Data Fig. 20.**
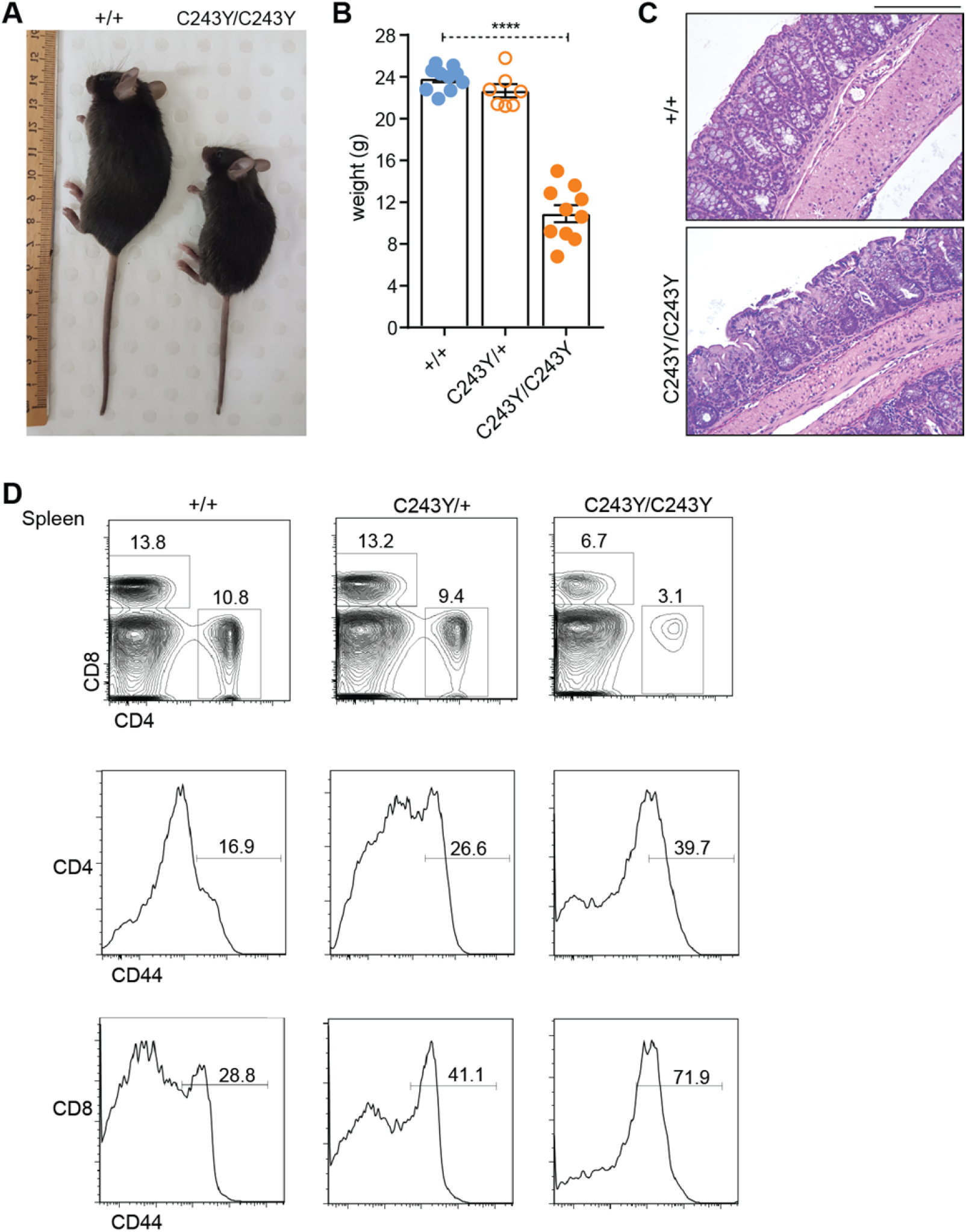
Histological analysis and immune phenotyping of A20^C243Y^ mice. **(A)** Representative photo of 8 week-old female *Tnfaip3*^+/+^ and *Tnfaip3*^C243Y/C243Y^ mice. Statistical analysis by one-way ANOVA: \*\*\*\**P* < 0.0001. **(B)** Weights (g) 8 week-old male *Tnfaip3*^+/+^, *Tnfaip3*^C243Y/C243Y^ and *Tnfaip3*^C243Y/C243Y^ mice. **(C)** Hematoxylin and eosin stained sections of colon from representative female mice in (A) (scale = 200 µm). **(D)** Flow cytometry of the percent CD4^+^ and CD8^+^ splenocytes (top panel) and the percentage of each subset that are CD44^hi^ activated/memory cells (histograms).

**Extended Data Table 1.** Allele frequencies of the Denisovan T108A;I207L haplotype in the Simons Genome Diversity Project^22^, as presented in (Extended Data Fig. 2B).

| **Region** | **Denisovan**<br>***TNFAIP3***<br>**alleles** | **Total**<br>***TNFAIP3***<br>**alleles** | **Denisovan**<br>**allele**<br>**frequency** |
| --- | --- | --- | --- |
| Africa | 0 | 88 | 0.00 |
| West Eurasia | 0 | 150 | 0.00 |
| Central Asia | 0 | 54 | 0.00 |
| South Asia | 0 | 78 | 0.00 |
| East Asia | 0 | 94 | 0.00 |
| Americas | 0 | 44 | 0.00 |
| Oceania & Island SE Asia: | 23 | 50 | 0.46 |
| (Brunei) | 1 | 4 | 0.25 |
| (Australia) | 1 | 4 | 0.25 |
| (Papua New Guinea) | 19 | 30 | 0.63 |
| (Bougainville) | 1 | 4 | 0.25 |
| (New Zealand) | 1 | 2 | 0.50 |

**Extended Data Table 2.** Polymorphic observed (black) and imputed (red) variants within a putatively adaptively introgressed haplotype in Melanesians (6:138160925-138246514, ^29^). Ancestral, modern human (b37), Denisovan, and Altai Neanderthal genotypes are shown at each position for comparison. Genomic coordinates are based upon the hg19 reference.

| Chr | Position | rsID | Missense | Ancestral | b37 | Denisovan | Altai Neanderthal |
| --- | --- | --- | --- | --- | --- | --- | --- |
| 6 | 138161013 | rs6918329 |  | G | A | G | G |
| 6 | 138174328 | rs2027276 |  | C | A | A | C |
| 6 | 138190154 | rs3757173 |  | G | A | A | G |
| 6 | 138195402 | rs596493 |  | C | T | C | C |
| 6 | 138196008 | rs376205580 | T108A | A | A | G | A |
| 6 | 138196957 | rs141807543 | I207L | A | A | C | A |
| 6 | 138223082 | rs17066926 |  | C | T | C | C |
| 6 | 138241185 | rs4895499 |  | G | A | G | G |

**Extended Data Table 3.**
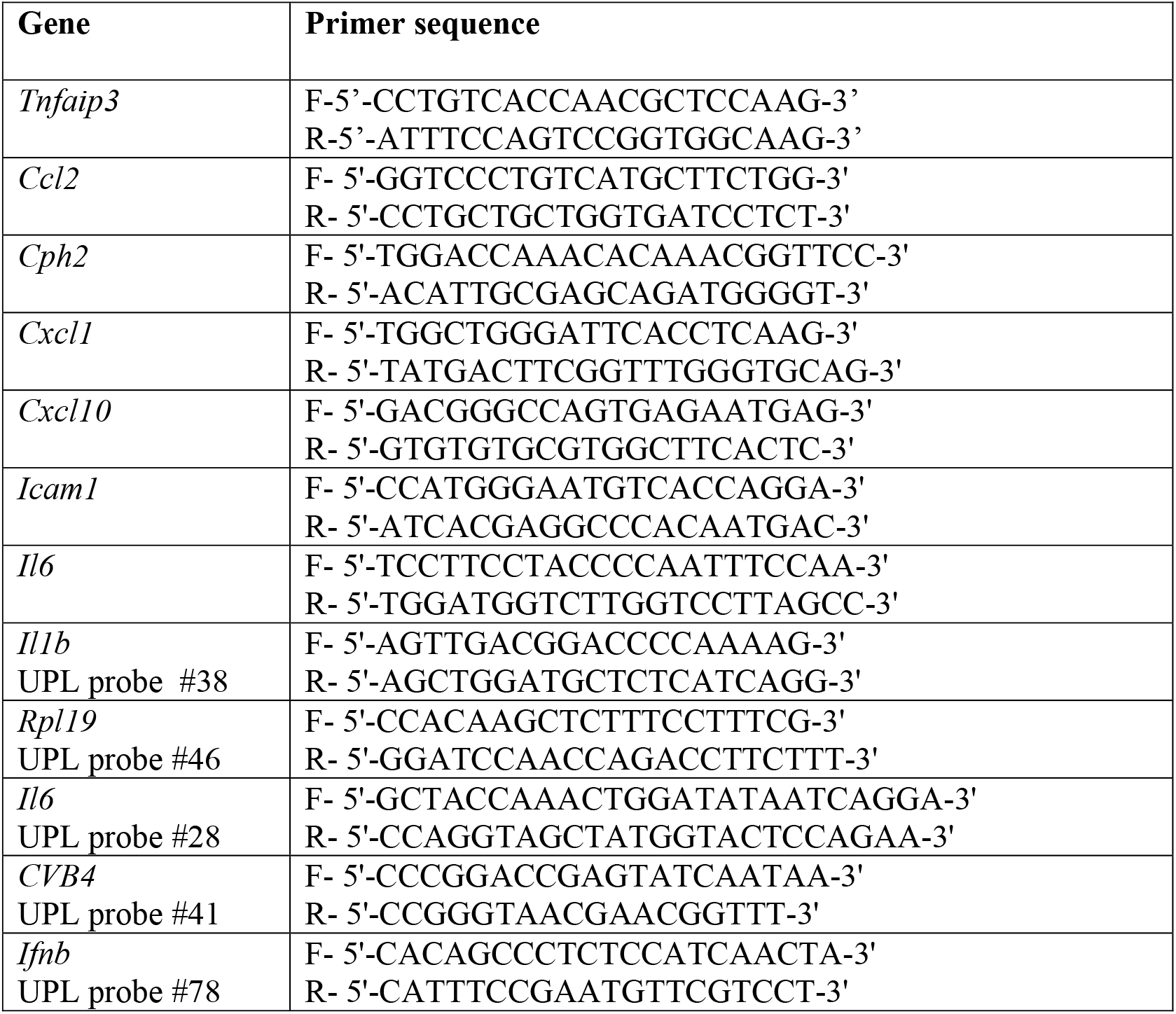
Mouse primers used for qRT-PCR analysis of genes. Designed using Primer3 with sequences obtained from GenBank and synthesised by Sigma Aldrich (Australia). Primers from Roche Universal ProbeLibrary (UPL) System Assay are indicated.

**Extended Data Table 4.**
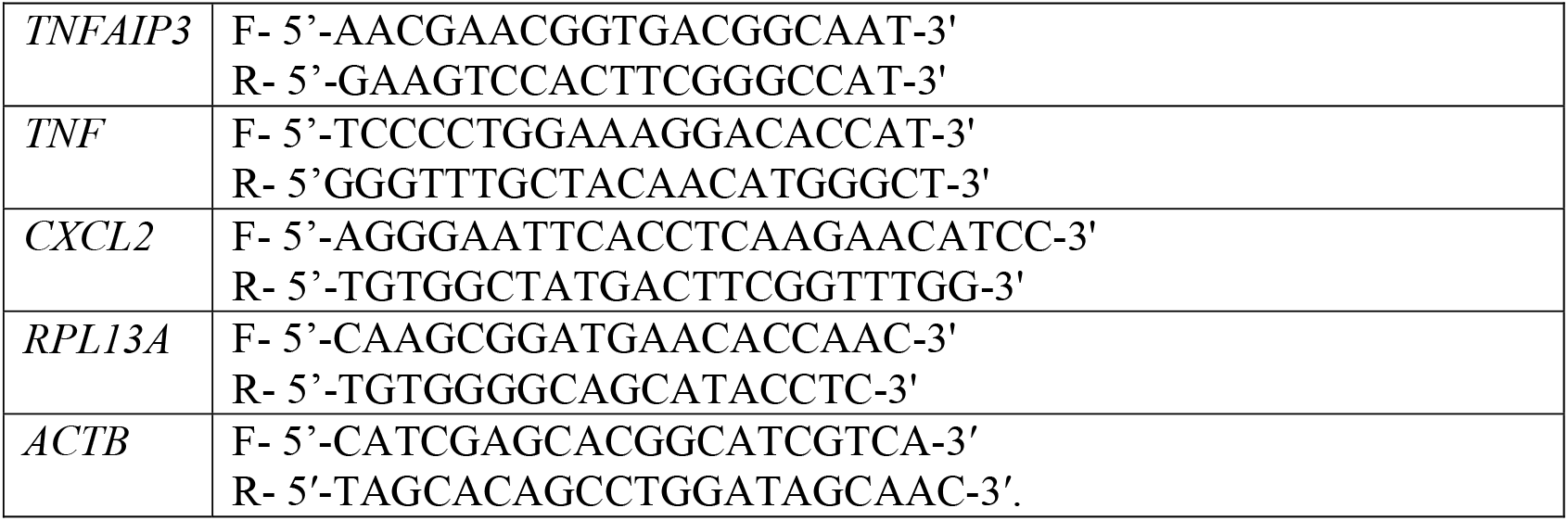
Human primers used for qRT-PCR analysis of genes. Designed using Primer3with sequences obtained from GenBank and synthesised by Sigma Aldrich (Australia).

**Extended Data Table 5.**
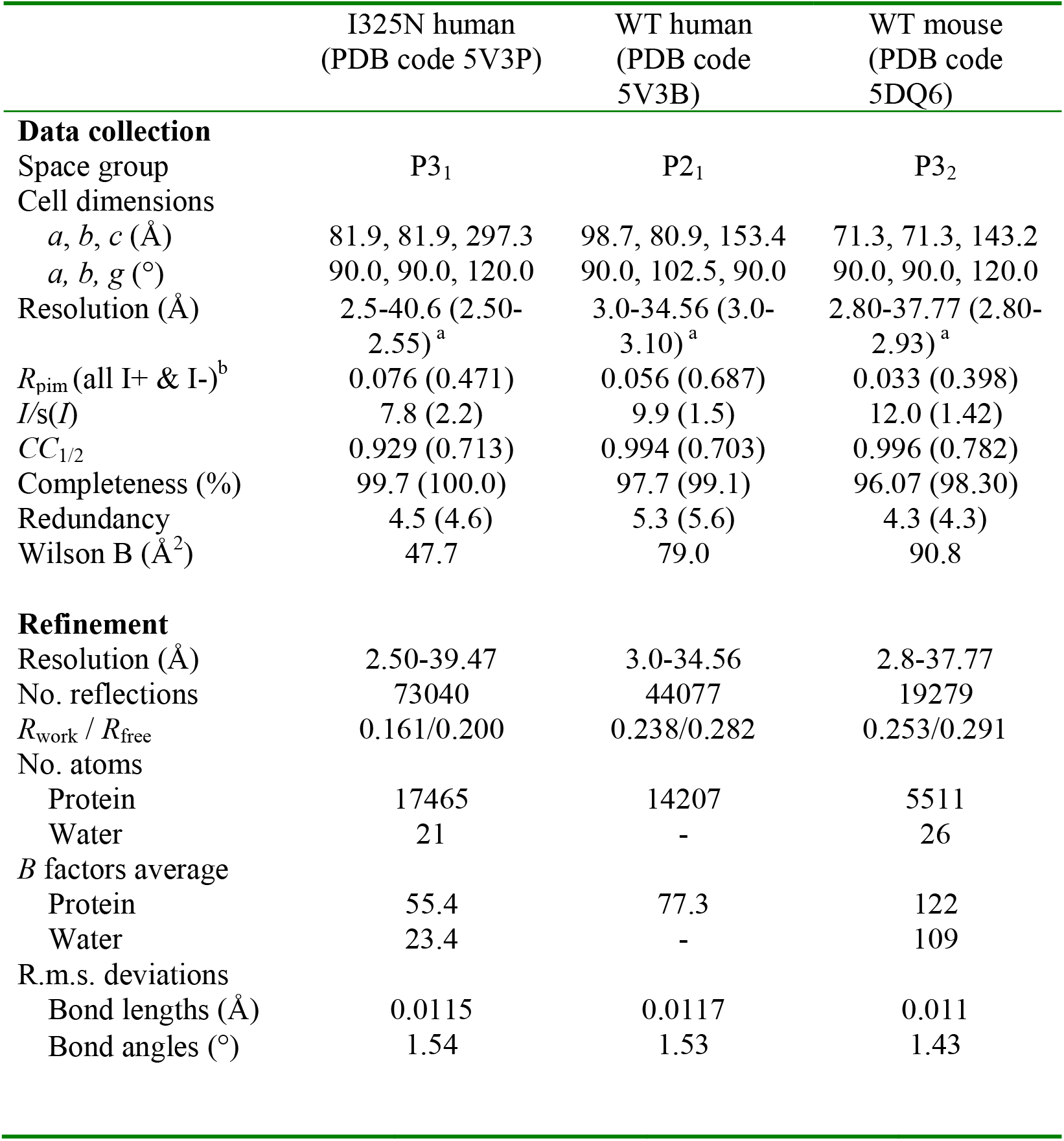
Data collection and refinement statistics (molecular replacement).

